# Species-specific drivers of genetic diversity are decoupled from landscape connectivity and plant community diversity

**DOI:** 10.64898/2026.06.25.734591

**Authors:** Lina Abdelwahed, Lisa Favre-Bac, Neda Rahnamae, Freya Way, Noémie Poulain, Tahir Ali, Anu Eskelinen, Irène Till-Bottraud, Juliette de Meaux

## Abstract

Understanding how habitat connectivity shapes biodiversity remains a major ecological challenge. In particular, the roles of connectivity and ecological heterogeneity in co-variation between plant species diversity and intraspecific genetic diversity are not well understood. We combined species distribution modelling, resistance-to-movement mapping, landscape connectivity analysis and population genomics to investigate diversity patterns in three wet-meadow herbs, *Scorzonera humilis, Oenanthe peucedanifolia* and *Lychnis flos-cuculi*, and their surrounding plant communities. Connectivity inferred from species distribution models was similar across species, but it did not correlate with plant community diversity. Some connectivity metrics explained genetic diversity, but only in *O. peucedanifolia*. Interestingly, environmental drivers of genetic differentiation were highly species-specific despite spatial co-occurrence. Furthermore, genetic diversity of two of the three species changed with the presence of some species in the community, but it was consistently unrelated to local plant community diversity. Overall, the processes shaping within-species biodiversity may differ fundamentally from those structuring metrics of habitat connectivity and plant species diversity, with important implications for conservation.

## Introduction

In the face of worldwide biodiversity loss, species persistence and the maintenance of diverse communities depend on how ecological features of the landscape, such as connectivity, can dampen the effect of the increased fragmentation caused by habitat destruction (Macarthur and Wilson 1967; Holyoak et al. 2005; Haddad et al. 2015; Damschen et al. 2019; IPBES 2019). The diversity of the species that compose specialized communities cannot be conserved without a clear understanding of how organisms move across heterogeneous landscapes (Brodie et al. 2025).

Connectivity impacts diversity at multiple levels and in two different ways. At the community level, high connectivity limits extinction by favoring recolonization but it may also decrease species diversity through promoting dominance of highly competitive species (Mouquet et al. 2002; Fahrig 2003; Matthies et al. 2004; Leibold and Chase 2017; Huovinen et al. 2025). Connectivity also impacts genetic diversity within populations in the same two ways. It can improve or decrease population fitness, depending on whether it introduces adaptive or maladapted alleles (Whitlock 2000; Lopez et al. 2009; González et al. 2020; Yeaman and Whitlock 2011; Andrade-Restrepo et al. 2019, Aitken et al. 2024). Genetic and community level diversities can also impact each other. Studies of experimental communities show that genetic variation can create eco-evolutionary feedbacks via alteration in species interactions and coexistence, and thereby alter the processes governing community assembly (Lankau and Strauss 2007; Des Roches et al. 2018; Wuest and Niklaus 2018; Frachon et al. 2019; Noto and Hughes 2020; Barbour et al. 2022). Yet, genetic diversity between populations is also shaped by the balance between gene flow and local adaptation, which respectively promotes or limits effective connectivity between populations (Yeaman, 2015).

Although landscape connectivity can impact both species and genetic diversity in the same ways, relationships between the two remain elusive (Whitlock 2014; Lamy et al. 2017). A key unresolved question is whether landscape connectivity will enhance both community and genetic diversity of species of the same community or whether genetic diversity follows species-specific patterns independent of the processes driving community diversity. Bridging this gap requires integrating landscape connectivity, population genomics, and community ecology within a comparative, multi-species framework, where these processes are compared across a common scale (Tack et al. 2012).

Recent advances in landscape modelling and high-throughput sequencing technologies now enable such analyses. Graph-based connectivity metrics based on species distribution models (SDM) allow spatially explicit quantification of potential dispersal pathways across heterogeneous landscapes (Urban and Keitt 2001; Duflot et al. 2018; Savary et al. 2024), while reduced-representation genomic approaches enable the characterization of genome-wide genetic variation in non-model species (Fitzpatrick and Keller 2015; Andrews et al. 2016). If species use the same dispersal corridors in a connected landscape, modelled connectivity should predict realized gene flow consistently across species.

Here, we integrate landscape ecology, community ecology and landscape genomics to quantify connectivity and its genetic consequences across three representative species within the same type of specialized and diverse communities. Focusing on wetland plant communities of the Massif Central (France), we combine habitat suitability models, resistance surfaces, graph-theoretical connectivity metrics, community composition data, and reduced-representation genome sequencing. Using three co-occurring wet-meadow species differing in dispersal characteristics (*Scorzonera humilis, Oenanthe peucedanifolia* and *Lychnis flos-cuculi*) as a model for genetic diversity in these communities, we ask: (1) do species of the same community share patterns of landscape connectivity, (2) are patterns of genetic diversity shaped by shared connectivity or by species-specific ecological drivers, and (3) does genetic diversity covary with community diversity, and if so, is this mediated by shared connectivity? By linking connectivity, genetic variation, and community structure across species, we show that even if species share broad patterns of connectivity, their genetic diversity responds to species-specific ecological drivers that are independent of those shaping community diversity.

## Material and methods

### Study species and sampling

We selected three species that characterize floodplain meadows within the Pilat Natural Park in France, are frequent enough to co-occur throughout the study area and are diploid (Fig S1-2, Suppl. Text). These species, *S. humilis*, *O. peucedanifolia* and *L. flos-cuculi* are all hermaphrodite perennials of meso-to hygrophilous meadows (Biere 1995; Colling et al. 2002). All three species are insect-pollinated with estimated seed dispersal distances of 15, 1500 and 1000 m, respectively (Wessinger 2021, Fig S2, Suppl. Text).

We sampled sites on three adjoining plateaus, Mornantais (7,825 ha), Pélussinois (12,662 ha) and Annonéen (18,052 ha). Further details on the species and sampled areas are given in supplementary information. In the 3 plateaus, plant populations of each species were sampled at 10 sites. For each population, the leaves of around 15 individuals were collected, preserved in paper envelopes and stored in glass containers with silica gel.

### Community level diversity

We conducted floristic surveys in the floodplain meadow plots where genetic material was sampled. In each area, ten 1 m² quadrats were positioned at randomly selected points. Within each quadrat, we recorded the presence of all plant species. We estimated species abundance within each plot as the relative frequency of occurrence of each species across the ten quadrats. This technique allows rapid completion of floristic surveys across larger areas. Based on these estimates of abundance data, we computed three *α*-diversity metrics per population: Species richness (number of species per area), the Shannon diversity index (*H′*), which accounts for both richness and evenness (Shannon 1948), and the Simpson diversity index (*1 – D*), which emphasizes species dominance (Simpson 1949). Diversity indices were computed using the *hill_taxa()* function from the *hillR R* package (Li 2018).

### DNA extraction and RAD-seq library Construction

Genomic DNA was isolated from ≥ 40 mg of silica-dried leaf tissue with the NucleoSpin 8 Plant II kit (Macherey-Nagel) and quantified with a Qubit BR assay as detailed in Suppl. Text. A total of 1,217 samples yielded sufficient DNA for further genomic analysis. Sixty-one RAD libraries, each pooling 20 barcoded individuals, were built following Etter et al. (2011) with the high-fidelity restriction enzyme *KpnI*-HF (NEB) and 6-bp inline barcodes (O’Leary et al. 2018). Libraries (30 µL) were sequenced at the Cologne Center for Genomics on an Illumina platform, sequencing was paired-end with 150bp reads, yielding ∼120 M reads per library. RADseq reads of *L. flos-cuculi, S. humilis* and *O. peucedanifolia* were mapped to the reference genomes of closely related species *Silene latifolia* (Moraga et al. 2025), *Avellara fistulosa* (ERGA, https://www.erga-biodiversity.eu/) and *Oenanthe javanica* (Feng et al. 2025), respectively. Sequencing data was processed as described in Suppl. Text.

### Genetic diversity

Genetic diversity within populations was quantified following standard methods (Suppl. Text). Population structure was assessed using both principal component analysis (PCA) and global ancestry inference. PCA was conducted using PLINK v1.9 (Chang et al. 2015), which converted the filtered VCF to binary format (*--make-bed*) and calculated principal components with the *-- pca* algorithm. The resulting eigenvectors and eigenvalues were imported into R v4.4 for visualization with *ggplot2* (Wickham 2016). Global ancestry was inferred using ADMIXTURE v1.3.0. (Alexander et al. 2009). Prior to running ADMIXTURE, the filtered SNP dataset was converted to PLINK format, and no further LD pruning was performed because for RAD-seq data the 300 bp thinning already reduced physical linkage to individual sequenced fragments. ADMIXTURE was run for K = 1–10 using the default block relaxation algorithm and five-fold cross-validation (*--cv*). The optimal number of ancestral clusters was determined by the minimum cross-validation error. For the *F_ST_* index of genetic differentiation between populations, the package *STAMPP* in R was used (Pembleton et al. 2013).

### Habitat suitability maps

We used 259, 472 and 537 presence points for *O. peucedanifolia, S. humilis* and *L. flos-cuculi,* respectively and used the *ENMEvaluate* function (ENMeval Package) and Maxent packages in R, along with 29 environmental variables to determine species distribution maps (Fig S3) (Muscarella et al. 2014; Phillips et al. 2017, Suppl. Text). With this, we created a map of favorable habitats using the *Maximum training sensitivity plus specificity Cloglog threshold* provided by Maxent (Phillips et al. 2017). This value is used to determine the cut-off value above which the probability of presence of a species defines the patch as either habitat or non-habitat (Liu et al. 2015). It was used in the *raster* calculator on *QGIS* (v. 3.34.4-Prizren, 2021) and provided a binary habitat suitability map for each species. We also transformed the species distribution map into a landscape resistance map representing the cost of movement in space using the formula of Keeley et al. (2016). The cost of movement ranges from 1 (probability of presence = 1) to 1000 (probability of presence = 0).

### Population connectivity metrics

We used Graphab version 2.8.6 (Foltête et al. 2021) to calculate connectivity metrics in the landscape. This landscape graph tool represents habitat patches as nodes connected by dispersion links that quantify the spatial heterogeneity of connectivity. Dispersion links are described along three dimensions: geographic distance, environmental distance and cost to movement. Graphab uses species distribution models (SDMs) as a data-driven and reproducible basis for quantifying the connectivity throughout the landscape (Stevenson-Holt et al. 2014; Duflot et al. 2018). The primary input data needed for Graphab are thus the habitat suitability maps and the landscape resistance maps, obtained as described above as well as a maximum dispersal distance (threshold) for each species. To incorporate uncertainties regarding short- and long-distance dispersal, we considered six dispersal thresholds: 500, 1000, 2000, 3000, 5000 and 10000 meters (Fig S4). For each dispersal threshold, three metrics were computed for each patch to reflect connectivity: the capacity of the patch, which reflects its ability to host a population, the Betweenness centrality (BC), which reflects whether a patch lies on movement pathways and can act as a stepping stone for dispersal (Boulanger et al. 2020) and the flux (F), which measures how well a patch is connected to neighboring patches within the dispersal distance threshold. F thus reflects the existence of corridors in the landscape, while BC highlights important stepping stones in the landscape.

### Partial Least Square Regressions

To understand how habitat connectivity drives genetic diversity, we used Partial Least Square (PLS) Regressions from a custom script from Savary et al. (2022). The genetic diversity indices (*F_IS_* and *π*) were the response variables, whilst connectivity metrics Capacity, Flux (F) and Betweenness Centrality (BC) were the predictor variables. PLS regressions are an alternative to multiple linear regression and principal component regressions specifically designed for highly collinear explanatory variables and thus suited to test the effect of the numerous and often collinear connectivity metrics generated with a range of dispersal values. R^2^ quantifies the proportion of variance explained and Q^2^ the predictive ability of the model. A Q^2^ value of 0.098 and above indicates a significant predictive ability of connectivity metrics (Savary et al. 2022).

### Scales of genetic differentiation

To determine the spatial scale at which genetic differentiation is influenced by landscape features, we calculated the Distance of Maximum Correlation (DMC, van Strien et al. 2014) using the *graph4lg* package in R (Savary et al. 2021b). The DMC provides a specific distance threshold at which the correlation between genetic distance and geographic or cost distance is maximized. It pinpoints the precise spatial scale where landscape features most strongly influence genetic differentiation. For such approaches, the Euclidean genetic distance (*D_PS_*), which calculates genetic distance using Identity by state (IBS, Purcell et al. 2007, fewer shared alleles = higher distance), is more appropriate than *F_ST_* estimates because they reflect better contemporary landscapes (Savary et al. 2022). The DMC, which is in cost-distance units, was then converted to Euclidean distance in meters using the function convert_cd (R package graph4lg (Savary et al. 2021b)). To determine the maximum distance over which gene flow affects genetic differentiation, we computed genetic autocorrelation, which quantifies the decay of relatedness between individuals as a function of geographic distance. Because the distance at which kinship approaches zero reflects the spatial scale of historical gene flow, this analysis provides an estimate of the species’ dispersal capacity, independent of landscape characteristics. We calculated the mean kinship coefficient among individuals across 30 distance classes (in km) using SPAGeDi (Vekemans and Hardy 2004).

### Isolation by distance, by environment or by resistance to movement

In order to identify the features of landscape heterogeneity that best explain genetic diversity, we performed partial Mantel tests of association with each of three different ecological distance types: the Euclidean distance, the environmental distance, and the resistance distance representing the cost of movement. The Euclidean distance was computed using the *Distance matrix* tool in QGIS (v. 3.34.4, Prizren, 2021). We also computed an environmental dissimilarity matrix using the R package *raster* (Hijmans 2025), where the environmental values per coordinate of each sample were extracted. Environmental predictors (Table S2) were first screened for collinearity using pairwise correlations, and variables with *r* > 0.7 were not included together in subsequent analyses. The pairwise environmental distance was then calculated using the Gower dissimilarity index, a distance index that accommodates multivariate data of different types, implemented in the *daisy* function of the *cluster* package in R (Maechler et al. 2026). To compute the isolation due to the cost of movement, that is the accumulated cost along the path of least resistance between two points (Fig S5, Wang and Bradburd 2014; Herrera et al. 2017), we calculated the resistance distance with Graphab (v 2.8.6), using the landscape resistance map as input.

In order to identify the environmental factors that shape population differentiation, we used Generalized Dissimilarity Modelling (Ferrier et al. 2007; Freedman et al. 2010; Mokany et al. 2022). GDM discriminates the contribution of geographic (isolation by distance IBD) and environmental isolation (isolation by resistance IBR or by individual environmental factors) to genetic differentiation (F_ST_). GDM uses non-linear regression to identify predictors that significantly explain changes in allele frequency along measures of isolation by yielding I-splines (Glück et al. 2022). The maximum value of the curve quantifies the biological effect of the gradient while its shape reflects the changes in adaptive allele frequency as described in Fitzpatrick and Keller (2015).

As environmental factors, we used a set of climatic and soil variables and included local habitat characteristics, such as shade (derived from the local slope and orientation of the patch) or altitude, (see Table S3). We also included species abundances, and habitat patch capacity in this analysis, as indicators of variation in the composition of the community. We further incorporated species composition by extracting abundance data (0 to 10) for co-occurring plant species per site based on floristic surveys (species present in fewer than five sites were removed). To reduce multicollinearity, we computed Spearman correlation coefficients among variable pairs using the *cor()* function in R. We retained only one correlated variable from each species pair and abiotic variable pair for which |ρ| > 0.7. GDM models were fitted using 999 permutations to assess statistical significance of predictors using the function *gdm.varImp* (GDM, v 1.6.0-6, Glück et al. 2022).

## Results

### Species of the wetland meadow communities share similar corridors

Since the three species are characteristic species of the same wetland communities, we examined whether connectivity metrics correlated across species. The patch capacities, which reflect species abundances, were strongly correlated between *S. humilis* and *O. peucedanifolia* (Spearman r = 0.53, p<0.01), but independent from those of *L. flos-cuculi*. Connectivity metrics of flux were strongly correlated between species pairs, especially cost-based ones, indicating that they used similar connected corridors, but their betweenness centrality was independent, especially when considering cost to movement (Fig 1). We conclude that the overall habitat network serves the three species similarly in terms of accessible corridors, but species differ in the patches that serve as stepping stones.

**Figure 1.**
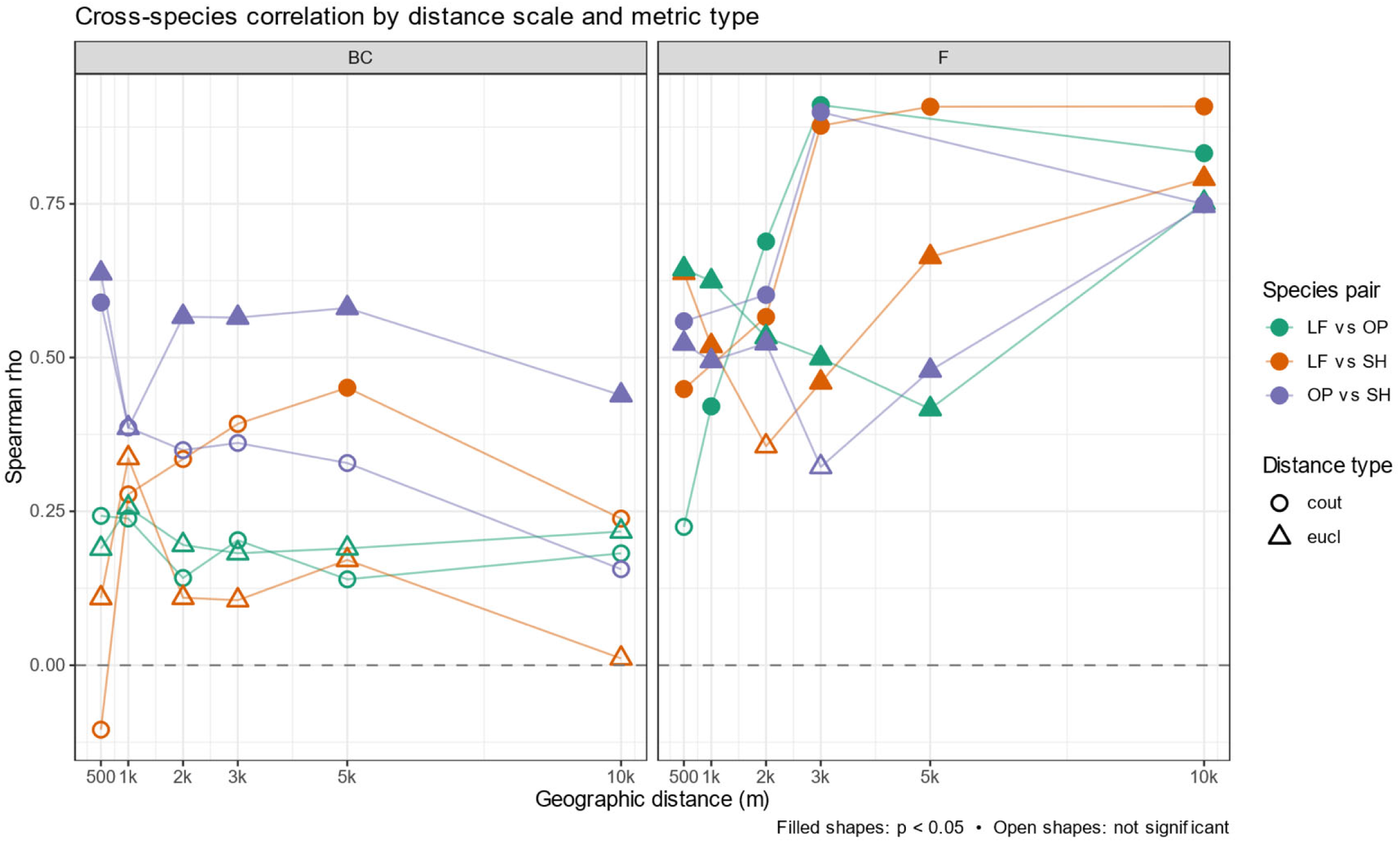
Pairwise correlation of connectivity metrics betweenness-centrality (BC) – left – and flux (F) – right – calculated for each species assuming increasing geographic distances of dispersal (500 m to 10 km). F measures how connected each population is, whereas BC reflects the populations that serve as stepping stones connecting populations. Green, orange, purple: correlation coefficient for connectivity metrics of *L. flos-cuculi* vs. *O. peucedanifolia*, *L. flos-cuculi* vs. *S. humilis* and *O. peucedanifolia vs. S. humilis,* respectively. Non-significant correlations have open shapes. Spearman correlation coefficient r was used to quantify and test for correlations

### RADseq data reveal different population structure for each species

With an average of 6 M sequencing reads per individual, we genotyped a total of 206 *S. humilis* individuals for 2886 single-nucleotide polymorphisms (SNP), 193 *O. peucedanifolia* individuals for 4452 SNPs and 352 *L. flos-cuculi* individuals for 5509 SNPs (Table S1, Fig S6). Species differed in their genetic structure. Using ADMIXTURE, *S. humilis, O. peucedanifolia* and *L. flos-cuculi* were arranged into 2, 3 and 4 ancestry groups, respectively, that were distributed differently among the plateaus (Fig 2, Fig S7-9). Genetic diversity within population was measured as the excess of heterozygotes (*Fis*) and the average number of pairwise differences among polymorphic sites (*π*). *Fis* values ranged from −0.14 to −0.06, −0.016 to 0.03 and −0.04 to 0.007 in *S. humilis, O. peucedanifolia, L. flos-cuculi*, respectively. Nucleotide diversity, estimated from both variant and invariant sites, ranged from 0.034 to 0.038 for *S. humilis*, 0.013 to 0.018 for *O. peucedanifolia*, and 0.037 to 0.046 for *L. flos-cuculi* (Table S4), respectively. *O. peucedanifolia* had the lowest nucleotide diversity, while *L. flos-cuculi* and *S. humilis* showed comparably high levels of genetic diversity, in agreement with known estimate for outcrossing plant species (Leffler et al. 2012; Ellegren and Galtier 2016; Chen et al. 2017). These estimates, however, should be taken with caution because genetic diversity estimates derived from RAD-seq are less accurate than those derived from whole genome sequencing (Dittberner et al. 2019). Pairwise F_ST_ values between populations were visualized as heatmaps for each species (Fig 3). *O. peucedanifolia* (F_ST_ of 0 to 0.15) and *L. flos-cuculi* (F_ST_ of 0 to 0.06) showed the highest levels of inter-population differentiation, with two clearly delimited clusters broadly corresponding to the northern and southern plateaus, whereas *S. humilis* displayed comparatively more continuous F_ST_ values across populations.

**Figure 2.**
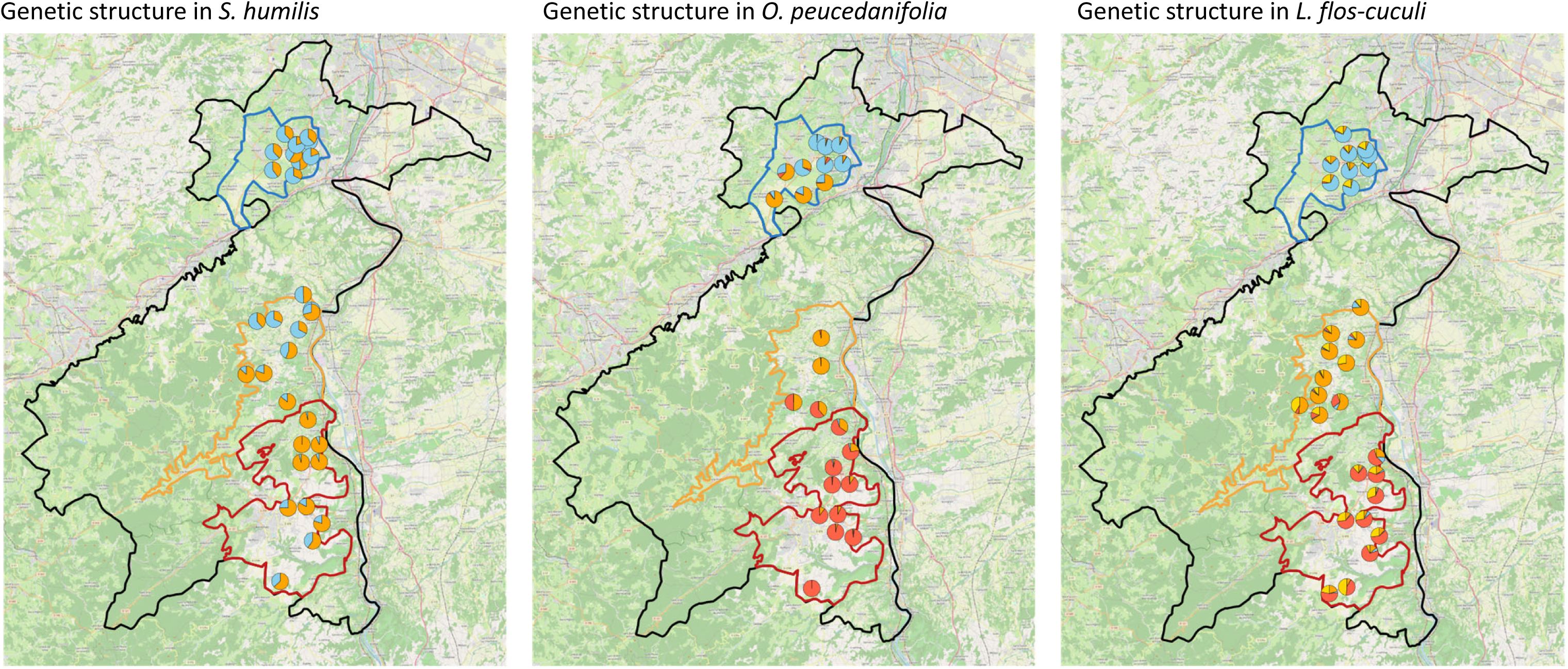
Geographic distribution of the genetic clustering of the three studied species, *S. humilis, O. peucedanifolia* and *L. flos-cuculi*.

**Figure 3.**
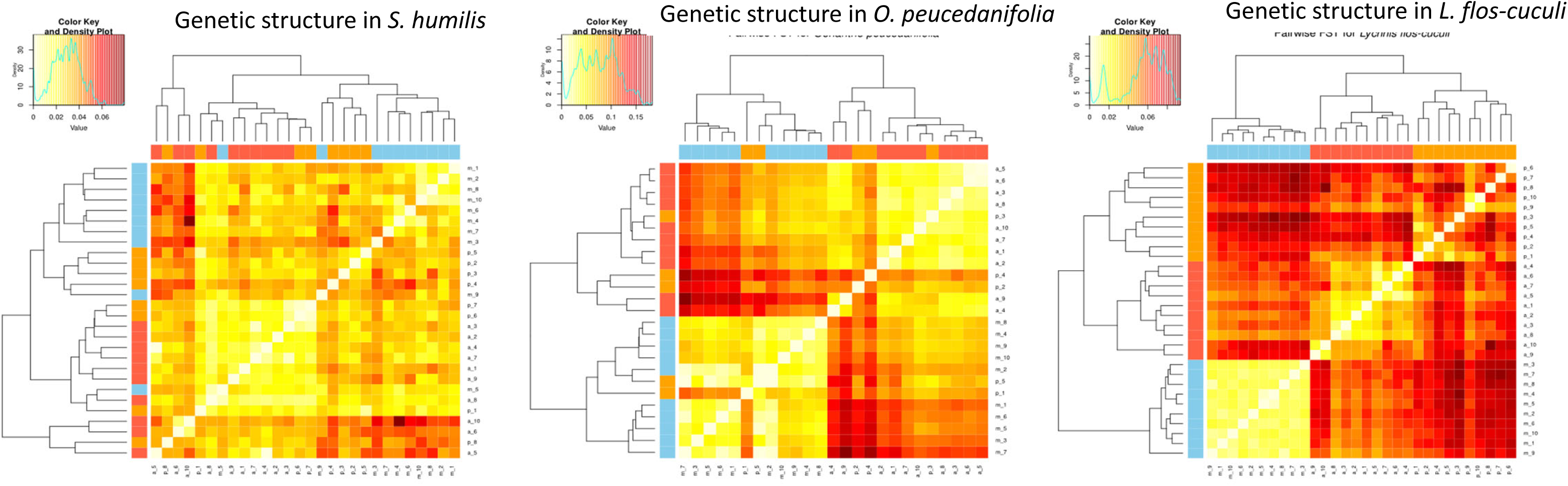
Pairwise linear F_ST_ heatmap of *S. humilis, O.peucedanifolia* and *L. flos-cuculi*. The heatmap color code represents pairwise F_ST_ considering different discrete F_ST_ bins, from low (orange and red) to high (bright yellow) genetic differentiation. The three plateaus are represented in blue (Mornantais), orange (Pelussinois) and red (Annonéen).

### Community diversity is unrelated to landscape connectivity of characteristic species

Because connectivity can shape diversity at the community level, we next tested whether the α-diversity of the surrounding plant community (species richness, Shannon index and Simpson index) was predicted by the connectivity metrics of each species, using Partial-least-square (PLS) regressions. For none of the three species did connectivity metrics predict community diversity: predictive ability Q² remained below the significance threshold of 0.098 in every case (PC1 predictive ability *S. humilis*, Q² = 0.02; *O. peucedanifolia*, Q² = 0.09; *L. flos-cuculi*, Q² = −0.08; Fig S11). These results indicate that habitat connectivity inferred from the presence and abundance of these three typical species of wetland meadows is unrelated to the processes that maintain species diversity at the community level.

### Genetic diversity of populations is sometimes associated with landscape connectivity

We further tested whether genetic diversity within the three focal species depended on their ecological connectivity. For *S. humilis* and *L. flos-cuculi*, connectivity metrics failed to predict genetic diversity (PC1 predictive ability Q² < 0.009, Fig S10). For *Oenanthe peucedanifolia*, however, the model predicting genetic diversity with connectivity metrics was informative (PC1 predictive ability Q² = 0.25; Fig 4). Genetic diversity within population was negatively correlated with the flux metric at high cost (17968 units of cost), but not with flux metric measured assuming lower cost of movements, Euclidean distance or betweenness measures. This result thus supports the notion that connectivity at long dispersal distances reduces genetic diversity in *O. peucedanifolia*, a pattern that may reflect founder events. PLS regression further showed that patch capacity was positively correlated with genetic diversity in this species, consistent with the idea that higher amount of available habitat associates with higher genetic diversity.

**Figure 4.**
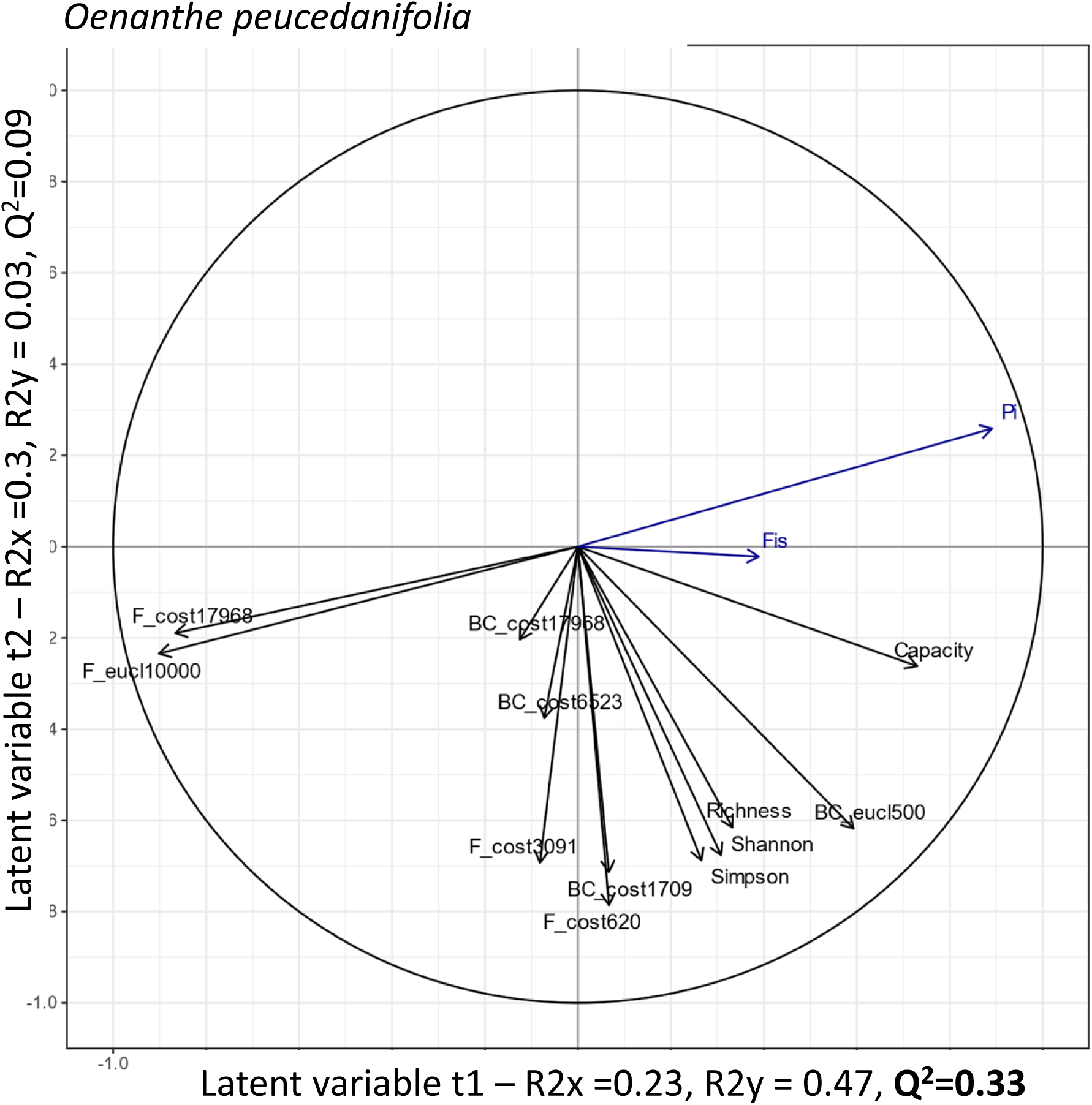
Predictive effect of species-specific connectivity metrics and community diversity on genetic diversity within populations of *O. peucedanifolia*. Partial least squares regression (PLS-R) biplot of *O. peucedanifolia* using connectivity metrics derived from the species’ own resistance map. Blue arrows = response variables; genetic indices (π, inbreeding coefficient Fᵢₛ); Black arrows = connectivity descriptors (patch capacity, betweenness centrality BC, flux F; Euclidean and cost, with dispersal thresholds indicated in subscripts) and α-diversity within patches (richness, Shannon, Simpson indices); The first two latent variables (t1, t2) are annotated with the proportion of variance they capture in X (R²X), in Y (R²Y) and with cross-validated predictive power (Q²). Q² values smaller than 0.1 indicate that the model has no predictive power and those greater than 0.5 indicate high predictive power. Acute angles denote positive associations, obtuse angles negative, and right angles negligible relationships. PLS-R models did not yield significant predictive power in the other species.

### Genetic similarity between populations confirms species differences in dispersal capacity

Genetic diversity can further inform about the dynamics of dispersal and gene flow. The geographic distance at which gene flow stops reflects the maximum dispersal capacity and can be estimated by the distance of maximum correlation (DMC) (van Strien et al. 2014). The DMC for *S. humilis, O. peucedanifolia* and *L. flos-cuculi* reached 500, 16700 and 10800 units of cost, translating to 45 m, 8496 m and 4331 m, respectively (Fig S12), showing that the distance at which gene flow stops impacting genetic relatedness differed markedly across species. These genetically based distances of dispersal aligned well with interspecies differences in physical dispersal capacities but indicated that dispersal capacities of the species are markedly greater than previously assumed. Indeed, physical dispersal capacities, were measured or inferred to reach 15, 1500 and 1000 m, for *S. humilis, O. peucedanifolia* and *L. flos-cuculi,* respectively (Vittoz and Engler 2007; Aavik et al. 2014).

Genetic autocorrelation describes how relatedness decays as a function of distance, and its distance limit reflects historical gene flow throughout the landscape (Anderson et al., 2010). Analysis of genetic autocorrelation revealed that differences between species in historical gene flow aligned with their dispersal kernels (Fig 5). In *O. peucedanifolia* and *L. flos-cuculi*, the kinship coefficient of genetic similarity declined steadily with distance and crossed zero at approximately 20 km and 10 km, respectively, indicating the spatial scale beyond which individuals are not more related than expected by chance. In *S. humilis*, kinship also declined at short distances but the correlogram was substantially noisier, with wide confidence intervals and a non-monotonic profile beyond the zero-crossing, so that no clear distance limit to relatedness could be defined.

**Figure 5.**
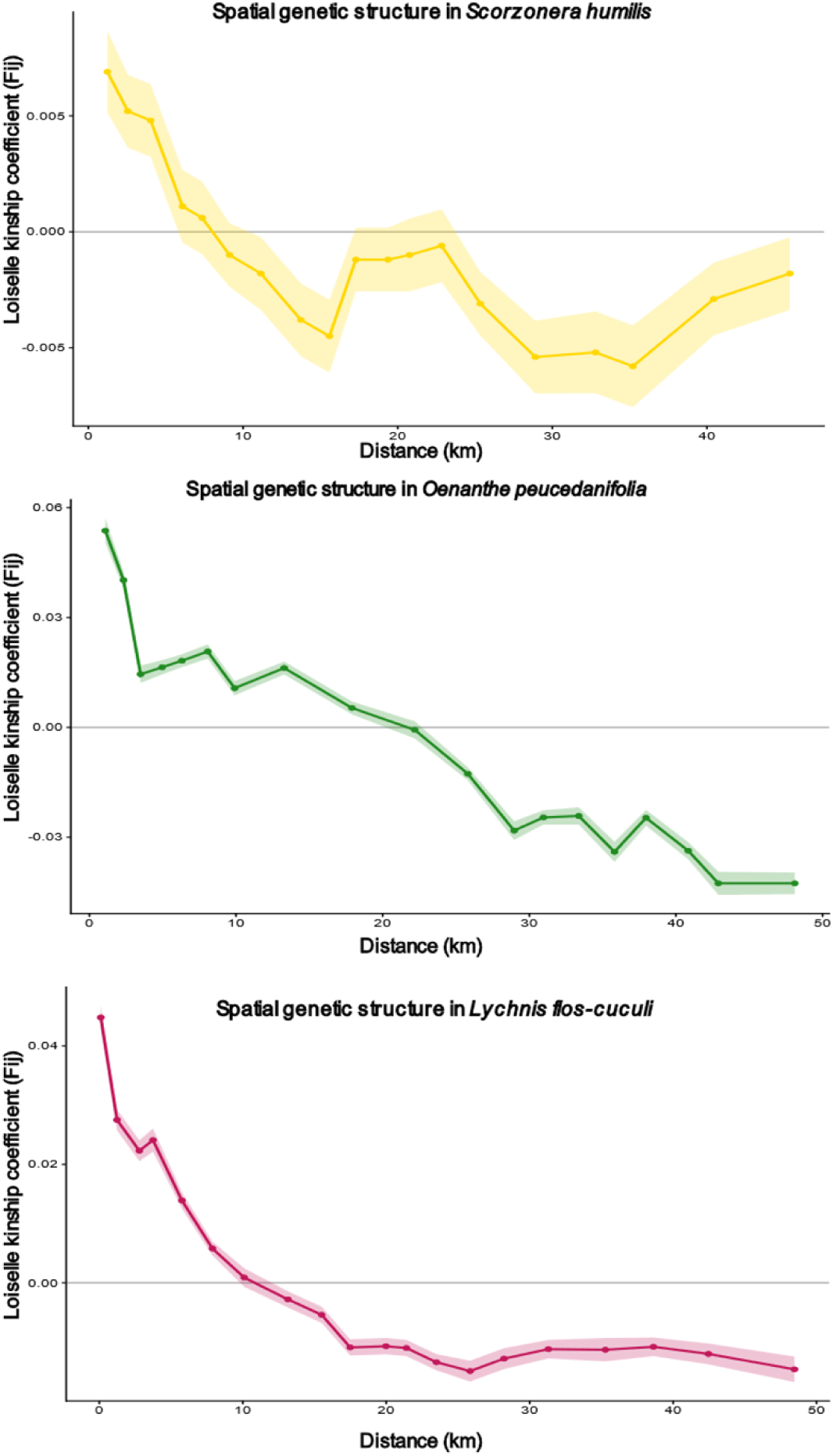
Autocorrelogram of the pairwise kinship coefficient (F₍ᵢⱼ₎) as a function of 30 classes of geographic distance between individuals in three different species (yellow=*S. humilis*, Green=*O. peucedanifolia*, Pink=*L. flos-cuculi*. Solid lines show the observed F₍ᵢⱼ₎ values for each distance class, and the lighter shaded envelopes of the same color represent the 95% confidence interval of F₍ᵢⱼ₎ under the null hypothesis of a random spatial distribution of genotypes.

### Species differ in how ecological distance explains genetic distance

The three species differed in the distance measure that best explained genetic distance (Table 1). For *S. humilis*, partial Mantel tests showed that both IBD (Euclidean distance) and IBR (resistance distance) played a role, but it was IBR that best explained differentiation between populations (Mantel coefficient = 0.196, 0.03, Table 1). For *O. peucedanifolia,* all three measures of distance had an effect, but IBD best explained genetic differentiation (Table 1). For *L. flos-cuculi,* it was IBE that best explained genetic differentiation better than either IBD or IBR (Table 1). Therefore, the three species differ in the factors that shape their genetic structure.

**Table 1.** Mantel tests and partial Mantel tests testing the effect of ecological and geographical distance on genetic differentiation in *S.humilis, O. peucedanifolia and L. flos-cuculi*. The first row tests the association between the genetic distance and each different ecological distances (geo: geographic Euclidean distance (IBD), cost: cost distance of the least-cost-path – cost of movement- (IBR), env: Environmental dissimilarity (IBE)). The following rows report the partial Mantel test after accounting for the geographic Euclidean distance. In grey is the distance accounted for in the partial Mantel test. For *L. flos-cuculi*, we performed two more partial Mantel tests, one after accounting for either the effect of cost of movement or one after accounting for the effect of environmental distance.

| *Species* | *Scorzonera humilis* |  |  |
| --- | --- | --- | --- |
| Type of distance | Geo | Cost | env |
| Mantel coeff | 0.255 | 0.324 | 0.128 |
| p-value | **0.0001** | **0.0001** | 0.055 |
| Partial Mantel coeff |  | 0.1961 | -0.0203 |
| p-value |  | **0.0289** | 0.5734 |
| *Species* | *Oenanthe peucedanifolia* |  |  |
| Type of distance | Geo | Cost | env |
| Mantel coeff | 0.766 | 0.754 | 0.513 |
| p-value | **0.0001** | **0.0001** | **0.0001** |
| Partial Mantel coeff |  | 0.1028 | 0.07121 |
| p-value |  | 0.1008 | 0.2079 |
| *Species* | *Lychnis flos-cuculi* |  |  |
| Type of distance | Geo | Cost | env |
| Mantel coeff | 0.504 | 0.612 | 0.5066 |
| p-value | **0.0001** | **0.0001** | **0.0001** |
| Partial Mantel coeff |  | 0.5269 | 0.476 |
| p-value |  | **0.0001** | **0.0001** |
| Partial Mantel coeff |  |  | 0.3803 |
| p-value |  |  | **0.0001** |
| Partial Mantel coeff |  | 0.3453 |  |
| p-value |  | **0.0001** |  |

### Specific environmental parameters associate with genetic distance

To better understand these patterns, we investigated the ecological factors and community components that explain genetic dissimilarity among populations. For *O. peucedanifolia* the overall GDM model explains 85.6% of the variation in the data (p < 0.001, 954 fitted permutations out of 999, Fig S13). In agreement with the above, we found that the most significant predictor was Euclidian geographic distance (p = 0.006, predictor importance of 0.87). Yet, the analysis also highlighted the effect of specific abiotic and community characteristics such as shade (p = 0.049, predictor importance of 10.1), the abundance of pepper saxifrage (*Silaum silaus*, p = 0.003, predictor importance of 3.63), an indicator of natural moderately damp and non-fertilized meadows (Wittig et al. 2006) and abundance of water forget-me-not, an indicator of wetter meadows (*Myosotis scorpioides,* p = 0.015, predictor importance of 3.72, Erzfeld et al. 2024). Euclidean geographic distance showed a steadily increasing spline, consistent with isolation by distance. Shade exhibited a nonlinear response, indicating accelerated genetic turnover at higher differences in canopy cover. The flat spline for *Silaum silaus* indicates that the sheer presence of this species affects genetic structure in *O. peucedanifolia*. In contrast, the abundance of *Myosotis scorpioides* displayed a steep spline, suggesting that the increasing presence of this species is strongly associated with changes in allele frequencies. Interestingly, microclimatic variables did not associate significantly with allelic turnover.

The GDM for *L. flos-cuculi* explained 92.3% of genetic differentiation among populations (p < 0.001; 943 fitted permutations out of 999, Fig S14). As in the above, Euclidean geographic distance was the strongest predictor (importance = 5.48; p < 0.001), indicating pronounced isolation by distance. *L. flos-cuculi* variation, however, was also structured by a subset of abiotic environmental factors describing environmental humidity levels. Among environmental and community variables, Bio08 – temperature of the wettest quarter- (p = 0.001, predictor importance of 1.951 with a steep increase), and Bio18 (average monthly precipitation of the warmest quarter, p = 0.047, predictor importance of 0.54) along with topographic slope (p = 0.049, predictor importance of 1.02) indicated that topographic heterogeneity and the mean monthly precipitation of the warmest quarter contribute to population differentiation. Two co-occurring species also showed significant associations with genetic dissimilarity: tall fescue (*Schedonorus arundinaceus*; p = 0.049) and world caraway (*Trocdaris verticillatum*, p = 0.050). These two species are indicative of meadows with different nutrient levels (Ellenberg and Leuschner 2010). The spline shapes suggest nonlinear responses, with steep increases in genetic turnover at higher environmental and compositional differences.

Finally, for *S. humilis,* the species that showed the weakest population structure, we were unable to fit a GDM model to the data that would refine our understanding of the factors driving resistance to movement in this species. Although GDM showed that the genetic diversity of two of the species was associated with the presence of four species in the community, it yielded no indication that the abundance of the three focal species influenced each other’s genetic diversity.

## Discussion

By integrating genomic, community, and environmental data, we quantified ecological connectivity and genetic diversity in three characteristic wet-meadow forbs. Although the species occupy the same habitat network and share broadly similar ecological corridors, the drivers of gene flow differed markedly among species and were largely independent of community diversity.

### No single landscape scale relevant to all taxa

Understanding how species and genetic diversity may be connected requires that these levels of diversity are studied across the same scale (Tack et al. 2012). These three diploid species coexist in the same wet-meadow communities, which are structured by the same hydrological network. The analysis of landscape connectivity performed here indicates that they tend to share the same corridors, but the habitat patches that serve as stepping stones generally differ. Fragmentation therefore operates at distinct effective spatial scales depending on both dispersal capacity and ecological barriers, rather than at a single landscape scale relevant to all taxa (Savary et al. 2021a).

At the genetic level, the three species exhibited patterns of isolation by distance that aligned well with their differences in dispersal capacities, but they also differed in the factors that best explain these patterns. Genetic differentiation in *L. flos-cuculi* was best described by landscape resistance of the local abiotic environment to movement, as previously reported for another wet-meadow area in Switzerland (Aavik et al. 2014). In *O. peucedanifolia*, genetic differentiation between populations was best described by isolation by distance, and in *S. humilis*, population structure was best explained by resistance to movement. These contrasts support the idea that each species is shaped by different dimensions of the landscape.

### Species-specific factors structure genetic diversity

PLS and GDM analyses further revealed species differences in the environmental correlates of genetic turnover. In *L. flos-cuculi*, turnover was associated with humidity, slope, and indicators of nutrient availability, whereas in *O. peucedanifolia*, it was linked to shade and indicators of flooding regimes. In *S. humilis*, resistance-based distances explained residual genetic structure despite extensive ancestry sharing. The decoupling of geographically and environmentally driven genetic structure (isolation by distance versus isolation by environment) shows that environmental gradients can structure allele frequencies independently of corridor availability. GDM indeed captures both neutral (drift/dispersal-driven) and potentially adaptive (environmentally-driven) components of genetic turnover (Fitzpatrick and Keller 2015). Together, these findings indicate that geographic distance, landscape resistance, and environmental variation all contribute to gene flow, but their relative importance is species-dependent. Species, indeed, do not necessarily share the same pattern of gene flow. Similar species-specific responses have been reported in other plant systems, where habitat suitability, environmental variation, or dispersal vectors influence genetic connectivity differently among co-occurring taxa (Nevill et al. 2019; Da Silva et al. 2021; Dellinger et al. 2022).

The genetic diversity within populations, in contrast to genetic structure, responded to connectivity in one of the three species. Only *O. peucedanifolia* showed a relationship between within-population genetic diversity and connectivity metrics. Notably, connectivity at long dispersal distances was associated with reduced genetic diversity in *O. peucedanifolia*, illustrating how corridors that connect distant populations can favor founder effects. In the other two species, demographic history, extinction-colonization dynamics, and historical bottlenecks likely outweighed contemporary structural connectivity. This pattern is consistent with theoretical and empirical studies showing that genetic diversity within populations often responds more slowly to fragmentation than genetic differentiation (Keyghobadi et al. 2005; Mona et al. 2014) and may retain signatures of past rather than contemporary connectivity (Batalha-Filho et al. 2024; Wade et al. 2026).

### Genetic diversity is decoupled from community diversity, but genetic differentiation tracks community composition

The goal of this study was to determine whether analyses of landscape connectivity can recapitulate genetic and community diversity. Indeed, if landscape connectivity jointly maintained genetic and community diversity, genetic diversity within populations should covary with the diversity of the surrounding community, and both should be predicted by connectivity. In this study, neither was the case: there was no relationship between genetic diversity within populations and α diversity of the plant community, and plant community diversity was not associated with the inferred landscape connectivity of the three species. Such decoupling between species and genetic diversity echoes previous work showing that local species richness does not reliably predict interpopulation genetic diversity, and that species-genetic diversity correlations can range from positive to absent or even negative (Kahilainen et al. 2014; Lamy et al. 2017; Noto and Hughes 2020).

Nevertheless, genetic differentiation is not independent of the community here, because we could show that it tracks the composition of the community. Indeed, genetic differentiation of *O. peucedanifolia* populations was associated with different indicator species of wetness, whereas that of *L. flos-cuculi* populations was associated with indicator species of nutrient levels. Whether these associations are caused by the presence of these species, reflect local adaptation to the conditions that these species indicate, or result from shared dispersal constraints warrants further investigation. These results, however, highlight that neither summary metrics of species diversity nor measures of landscape connectivity can recapitulate all dimensions of biodiversity.

### Limits of SDM-inferred landscape connectivity

Genetic diversity of *S. humilis*, *O. peucedanifolia* and *L. flos-cuculi* is structured by elements of landscape connectivity, such as environmental or resistance gradients, but metrics of landscape connectivity did not show a strong explanatory power. Landscape connectivity may in fact imperfectly reflect the true connectivity of the wet-meadow landscape. Inference of landscape connectivity from SDM-based habitat networks and graph-based metrics has been generally applied to animals (Correa Ayram et al. 2016; Savary et al. 2021a; Ficetola et al. 2011). Applications to plant species remain comparatively less common, as spatial habitat networks might incompletely account for the specificities of seed and pollen dispersal. In plants, dispersal syndromes, pollination systems, and life histories jointly determine the scale and sensitivity of genetic connectivity (Ballesteros-Mejia et al. 2017; Sinclair et al. 2018; Nazareno et al. 2021). Directly measuring pollen and seed dispersal, or experimentally validating life history traits, however, remains challenging (Beckman and Sullivan 2023; Sullivan et al. 2026). Genetic analyses therefore provide an empirical way to evaluate whether predicted landscape connectivity translates into effective gene flow, but they do not by themselves reveal the underlying dispersal mechanisms, life-history traits, or establishment processes that ultimately determine effective gene flow and migration routes (Metzger and Décamps 1997; Aavik et al. 2014; Mueller et al. 2014).

In conclusion, this multi-species landscape genomic study shows that genetic diversity indicates paths of connectivity that no other dimension of biodiversity can reflect. The interplay between ecological and genetic connectivity in species co-occurring in the same community is complex and multifaceted. Managing endangered wet-meadow systems will require moving beyond single-metric notions of connectivity toward integrative frameworks that jointly consider habitat amount, spatial configuration, resistance, environment, and genetic and community composition (Keyghobadi et al. 2005; Nielsen et al. 2022; Van Moorter et al. 2023; Brodie et al. 2025).

## Authorship statement

LFB, ITB, AE and JDM designed and supervised the study. LFB and NP collected plant material and community surveys, LA, LFB collected genetic data and performed analyses. NR, FW and TA provided technical and analytical support. LA wrote the manuscript with input from all authors.

## Acknowledgement

We thank Ricardo Feuerer for help in script preparation. This work was supported by the Deutsche Forschungsgemeinschaft (DFG Programs ME2742/13-1, TRR341 and EXC 2048/1-390686111) The fieldwork and research conducted in France by the CBN Massif Central was supported by the French region Auvergne-Rhône-Alpes (AURA). We thank the Cologne Center for Genomics (CCG), the Regionales Rechenzentrum der Universität zu Köln (RRZK), the German Network for Bioinformatics Infrastructure (de.NBI) and N. Guillerme, N. Blanchin and M. Piroux from the Conservatoire Botanique National Massif Central for their support.

## Data Availability

Filtered vcf files, species abundances, species distribution models, connectivity metrics and detaileds scripts are available on a Github (https://github.com/labdlw/project_pilat) and will be published in an ARC on the NFDI-Plant storage platform. DOI is pending. Raw fastq files were uploaded on ENA (reference number pending).

