## Supplementary text and figures for "Species-specific drivers of genetic diversity are decoupled from landscape connectivity and plant community diversity"

Figure S1. Map of the three 3 study areas studied in France.

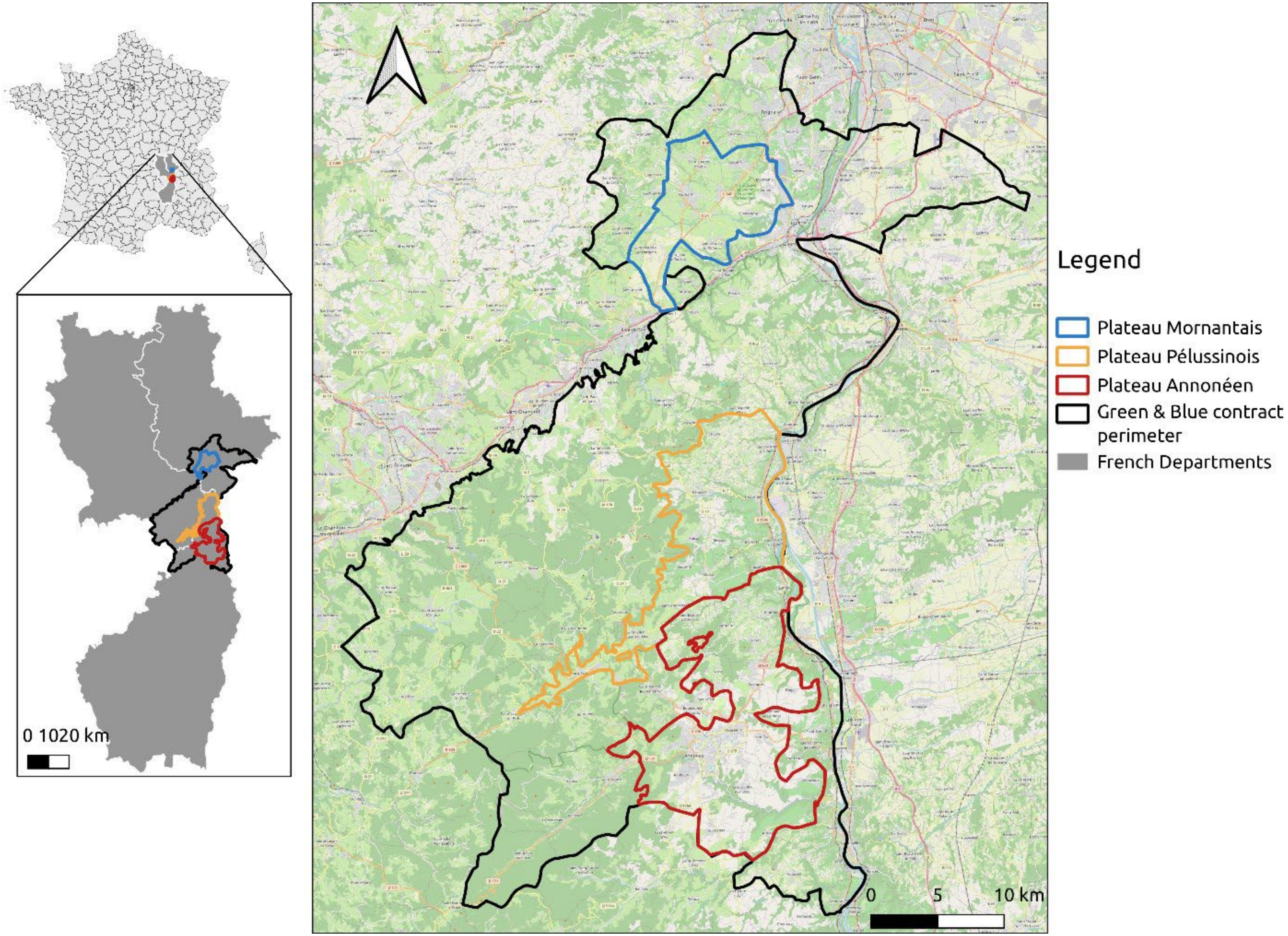

Figure S2. The inflorescence and the corresponding diaspore of each species, showcasing different dispersal mechanisms.

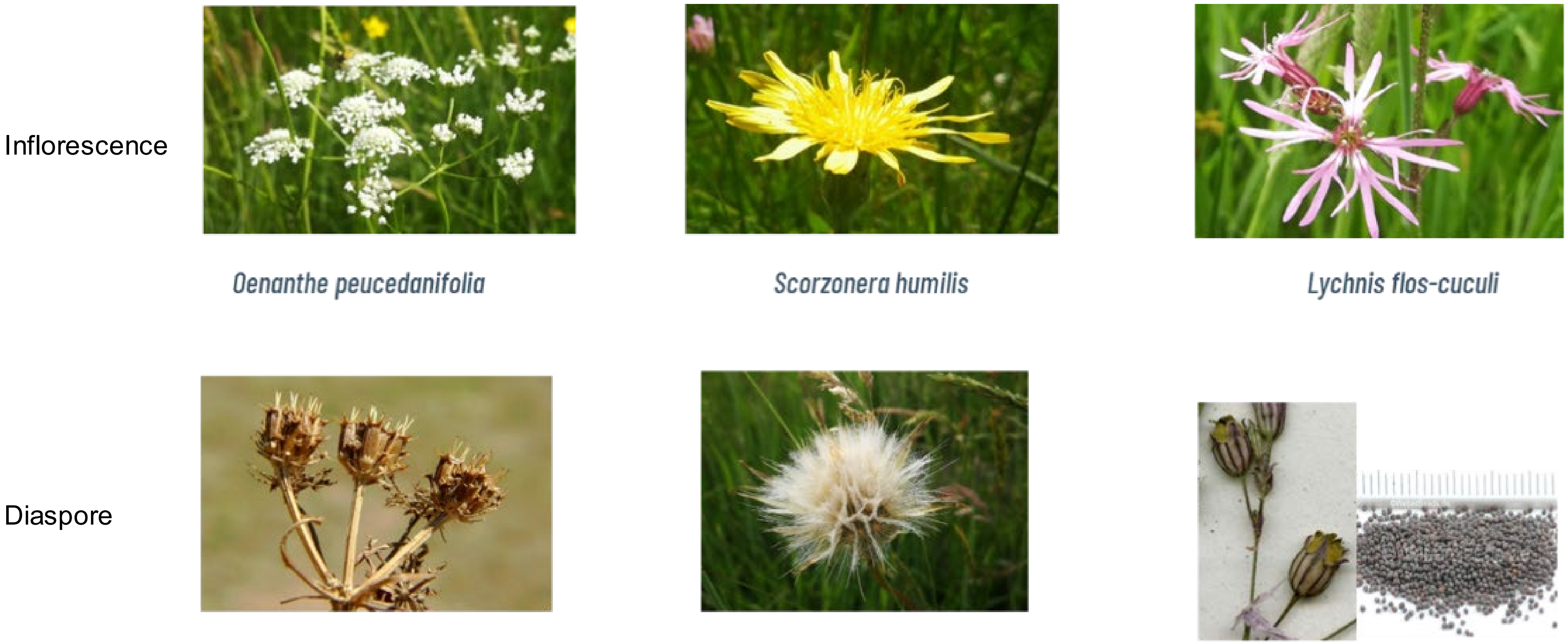

Figure S3. Construction of a floodplain meadow specialist map. Probability of presence of 5 species estimated through Species Distribution Modelling. A. *Scorzonera humilis. B. Oenanthe peucedanifolia.* C. *Lychnis flos-cuculi*. Dark blue colors correspond to a very low probability of presence (close to 0), Red colors correspond to higher probability of presence (close to 1).

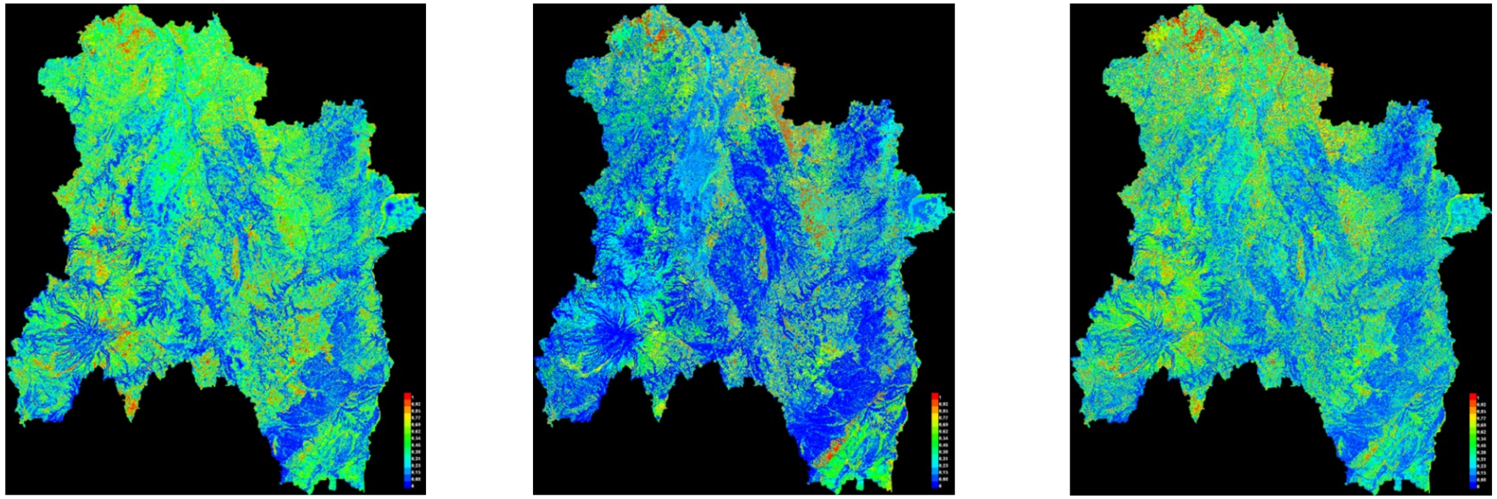

A. ***Scorzonera humilis*** **(AUC = 0.827)**

B. ***Oenanthe peucedanifolia*** **(AUC = 0.856)**

C. ***Lychnis flos-cuculi*** **(AUC = 0.798)**

Figure S4. An illustration of the different dispersal distances (meters) used to calculate connectivity measures. The higher the dispersal distance in meters considered, the more links (in black) are established between the habitat patches (in green).

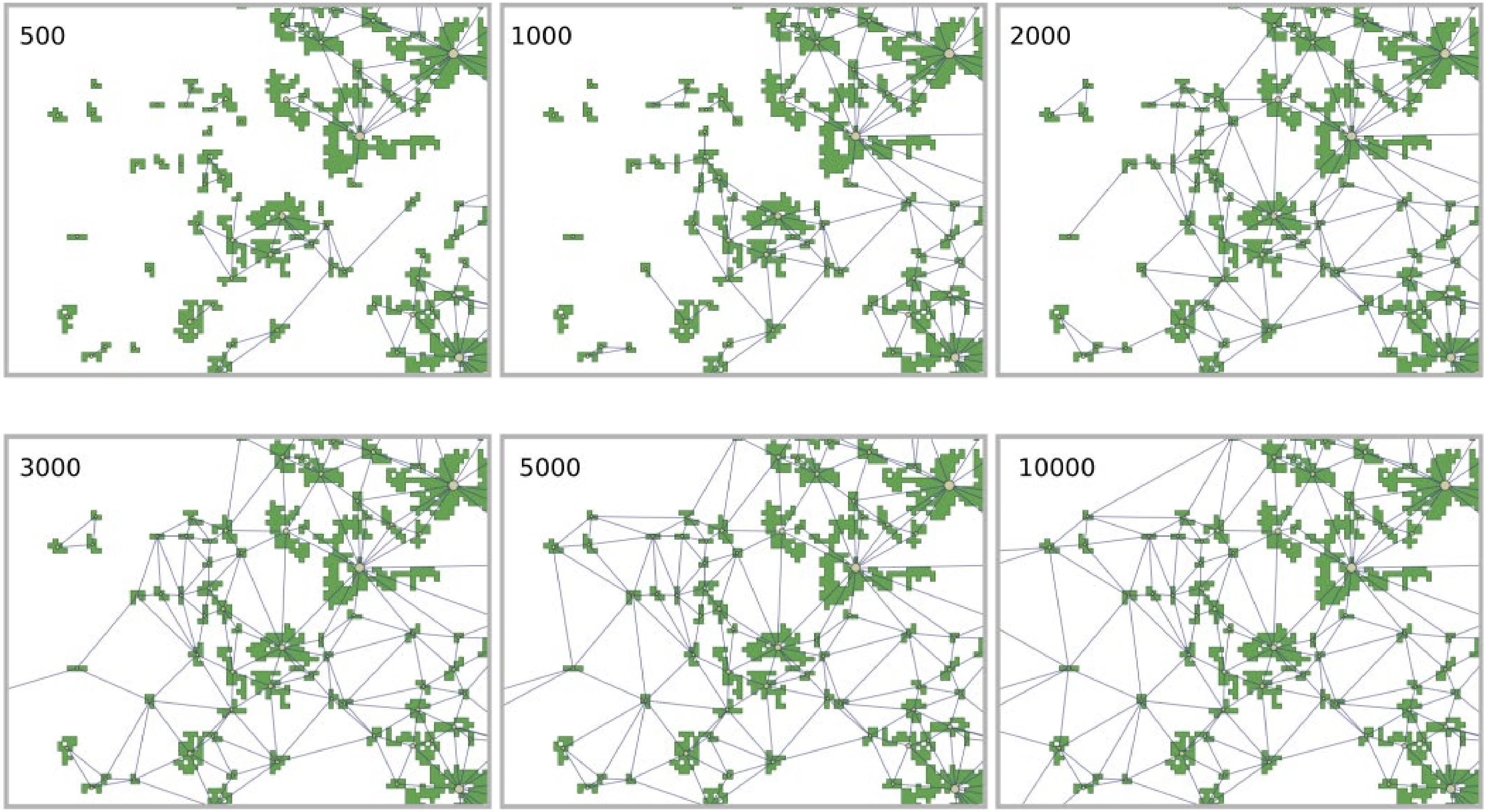

Figure S5. An illustration of habitat patches (in yellow) and the different ways they can be linked. Either through euclidean distance (A) or the least-cost path (B), where units of cost are measured through an accumulated cost throughout the path.

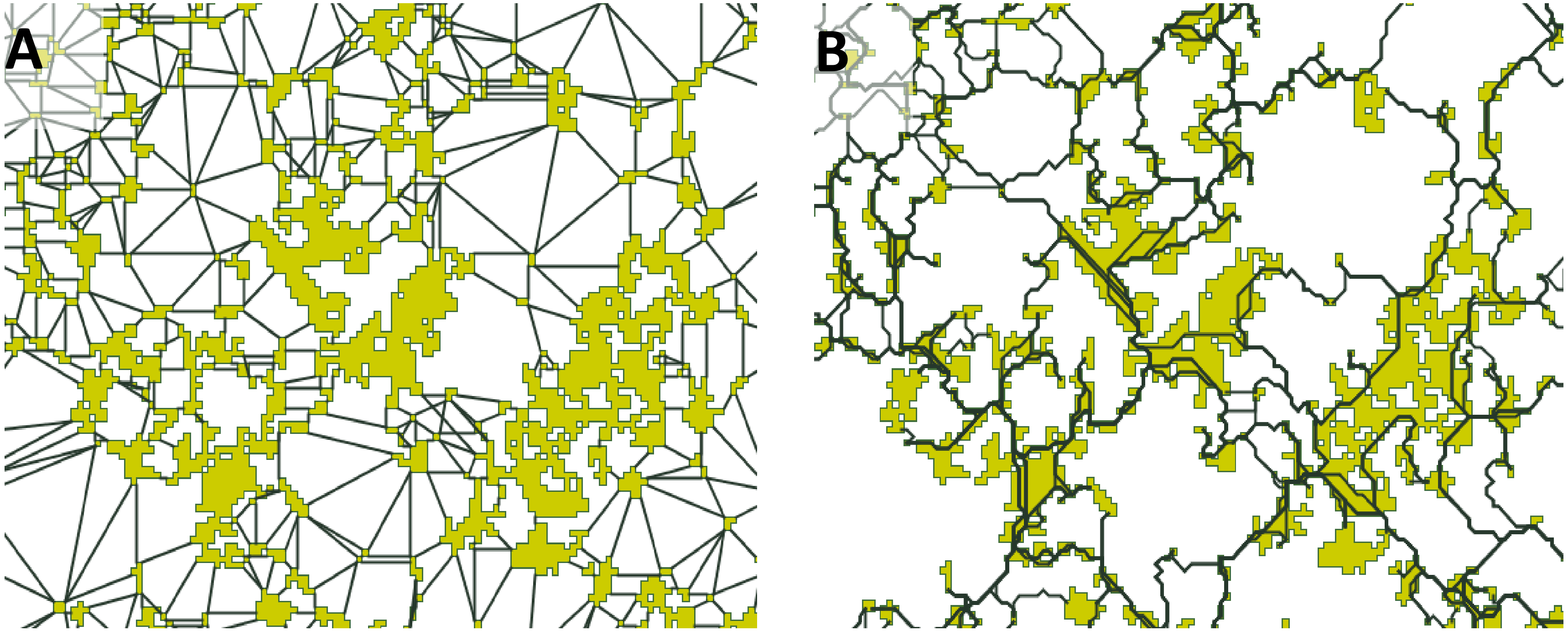

Figure S6. Number of retained reads (M) for all 1217 samples combined. The Red dashed line shows the median retained reads of 7M reads per sample.

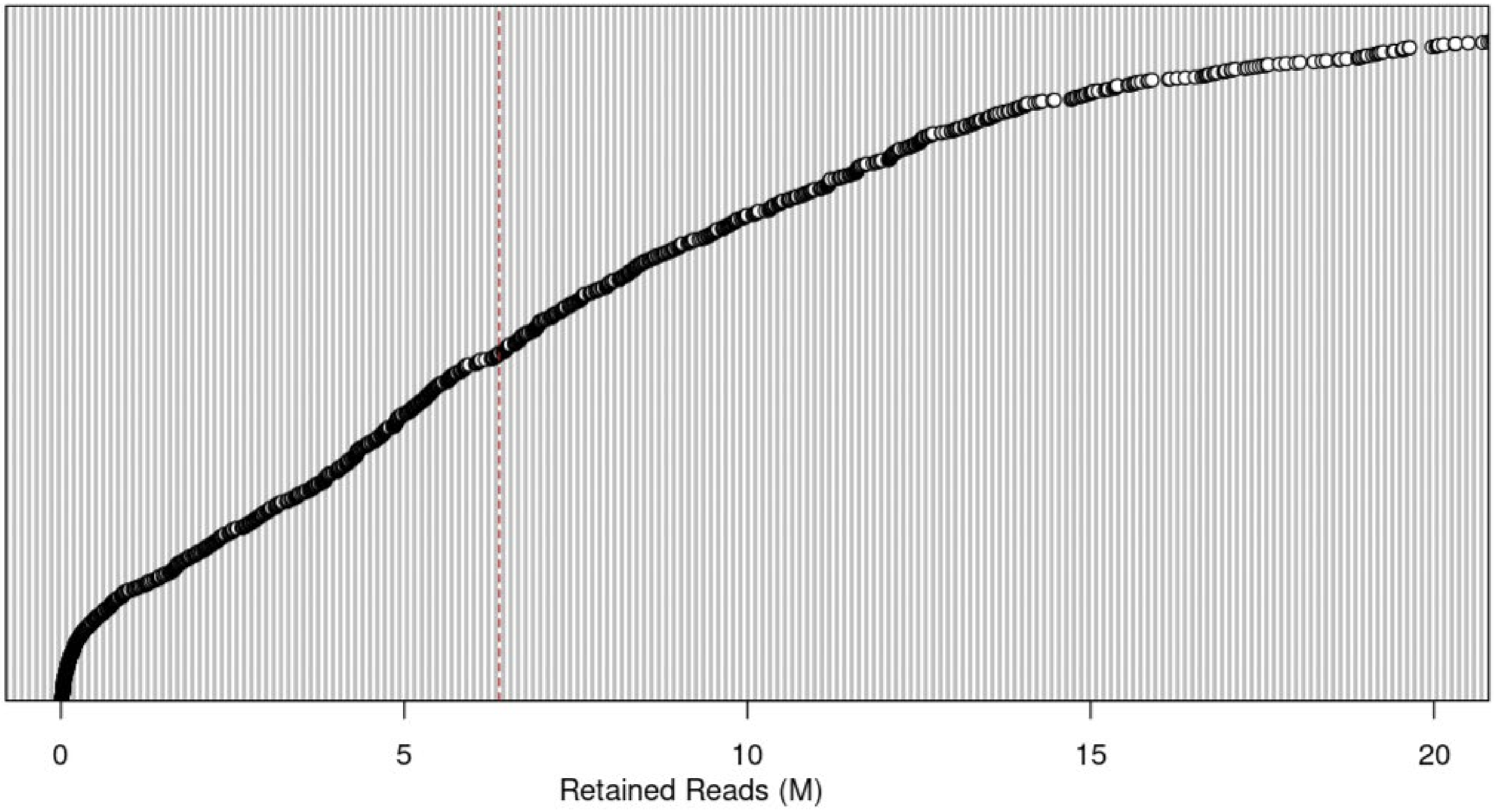

Figure S7.

ADMIXTURE was used to identify genetic groups within the population samples. *S. humilis* genetic diversity was best explained by two major clusters. Populations from the different plateaus are annotated as m (Mornantais), p (Pélussinois) and (Annonéen).

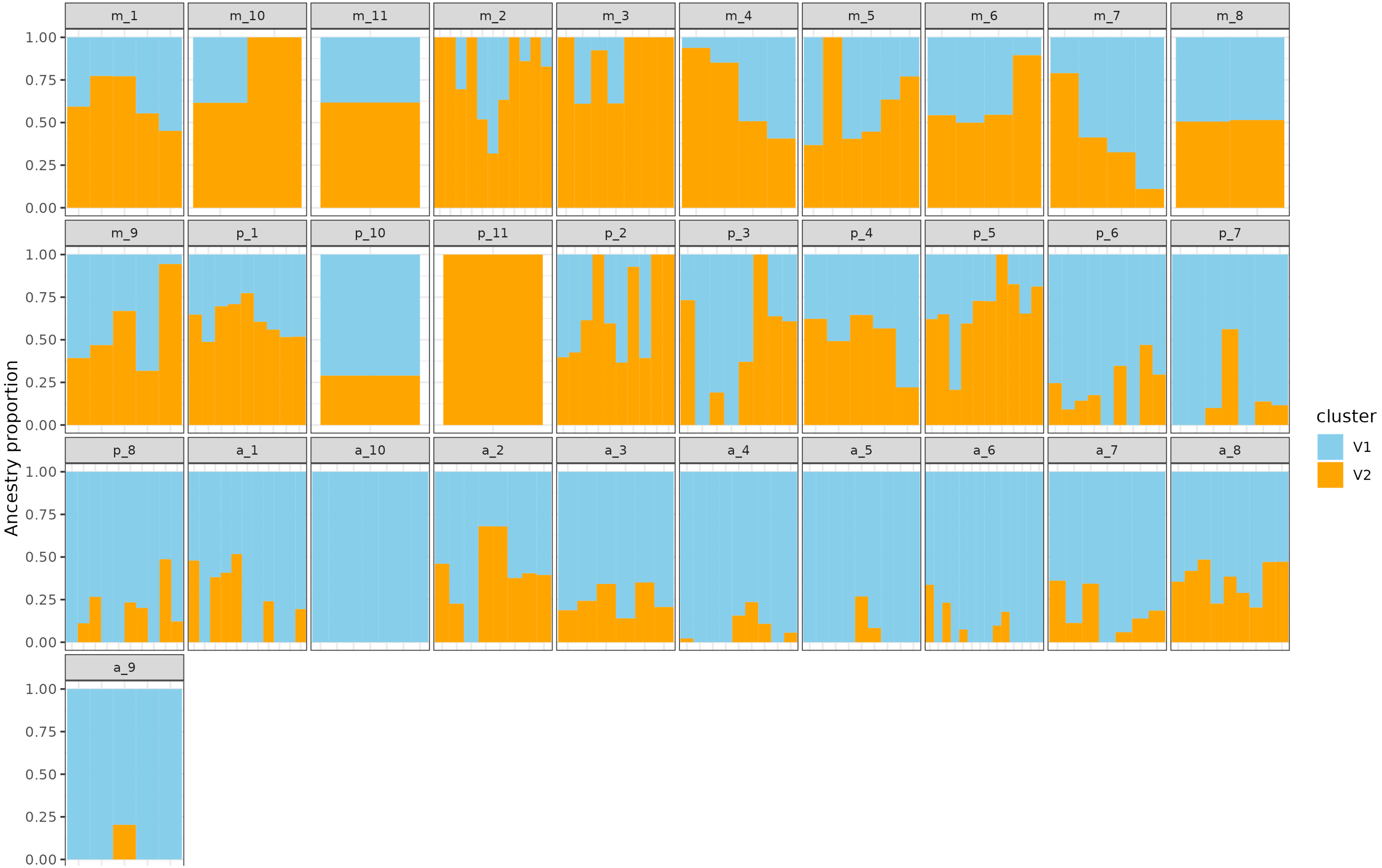

### Figure S8.

ADMIXTURE was used to identify genetic groups within the population samples. *O. peucedanifolia* genetic diversity was best explained by three major clusters. Populations from the different plateaus are annotated as m (Mornantais), p (Pélussinois) and (Annonéen).

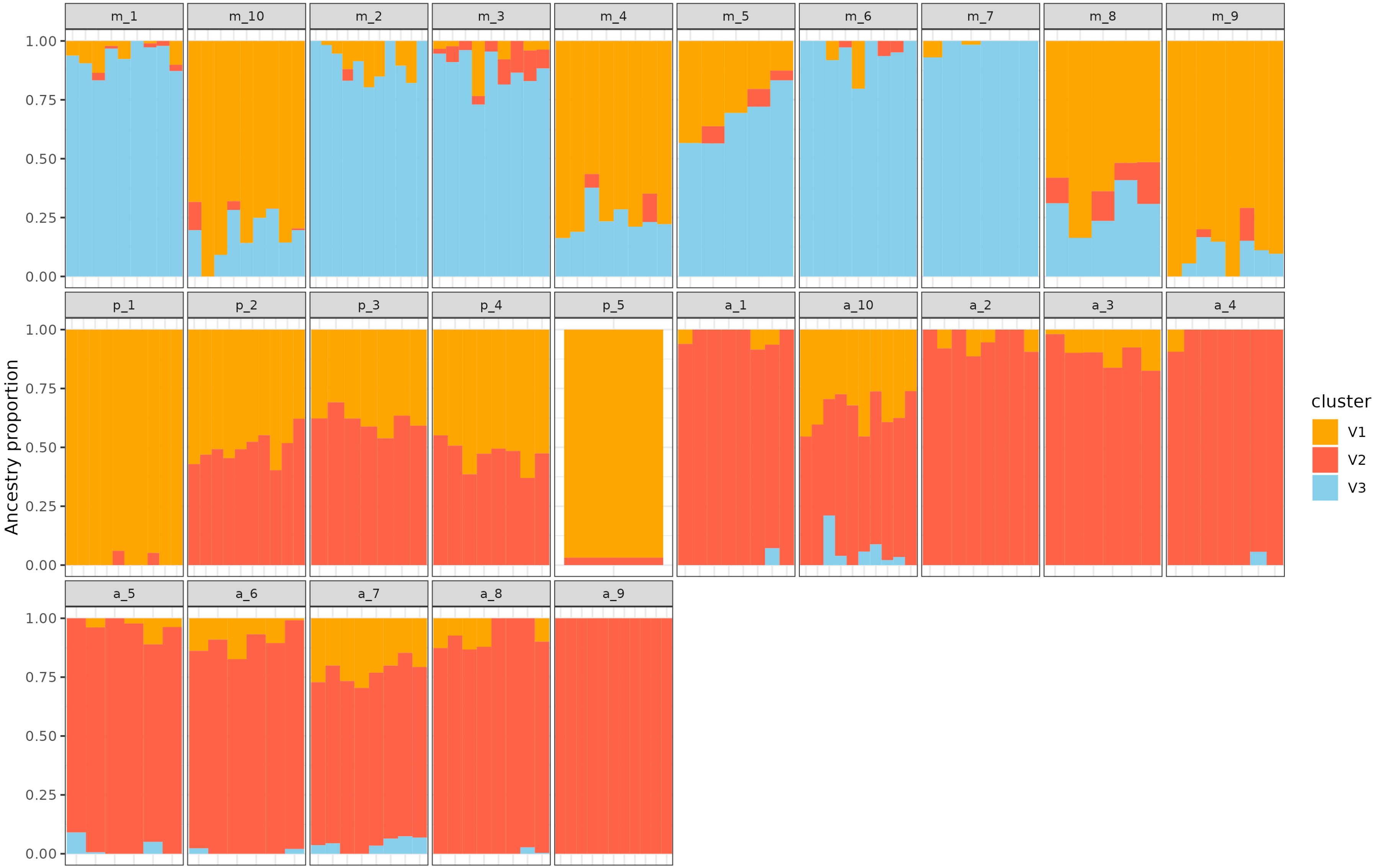

### Figure S9.

ADMIXTURE was used to identify genetic groups within the population samples. *L. flos.cuculi* genetic diversity was best explained by four major clusters. Populations from the different plateaus are annotated as m (Mornantais), p (Pélussinois) and (Annonéen).

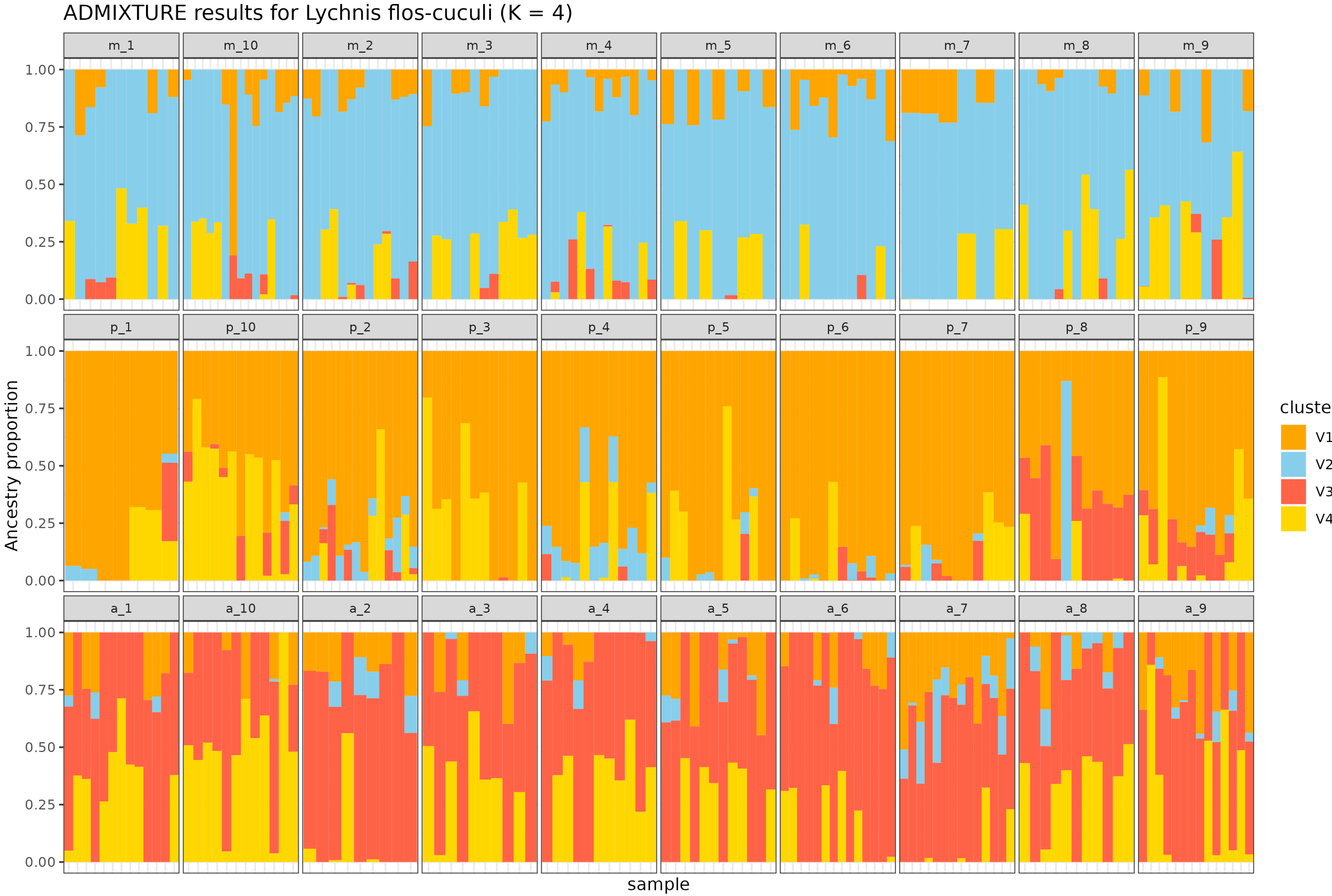

### Figure S10. Predictive effect of species-specific connectivity metrics & α-diversity  on genetic diversity within populations

Partial least squares regression (PLS-R) biplot of *Oenanthe peucedanifolia* using connectivity metrics derived from the species' own resistance map. Blue arrows = response variables ; genetic indices (π, inbreeding coefficient $F_{is}$); Black arrows = connectivity descriptors (patch capacity, betweenness centrality BC, flux F; Euclidean and cost, with dispersal thresholds indicated in subscripts) and  α-diversity within patches (richness, Shannon, Simpson indices); The first two latent variables (t1, t2) are annotated with the proportion of variance they capture in X ($R^2X$), in Y ($R^2Y$) and with cross-validated predictive power ($Q^2$). $Q^2$ values smaller than 0.098  indicate that the model has no predictive power and those greater than 0.5 indicate high predictive power. Acute angles denote positive associations, obtuse angles negative, and right angles negligible relationships.

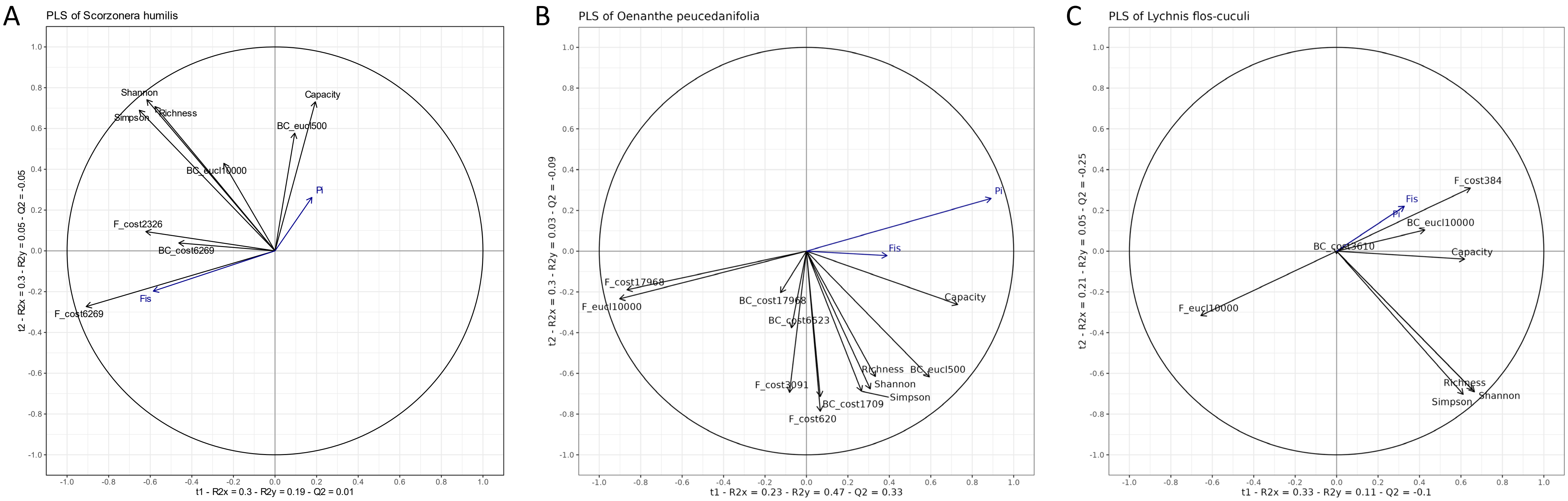

**Figure S11. Absence of predictive effect of species-specific connectivity metrics on plant community α-diversity.** Partial least squares regression (PLS-R) biplots of *Scorzonera humilis*, *Oenanthe peucedanifolia* and *Lychnis flos-cuculi*, each using connectivity metrics derived from the species' own resistance map. Blue arrows = response variables; α-diversity within patches (richness, Shannon, Simpson indices); Black arrows = connectivity descriptors (patch capacity, betweenness centrality BC, flux F; Euclidean and cost, with dispersal thresholds indicated in subscripts). The first two latent variables (t1, t2) are annotated with the proportion of variance they capture in X ($R^2X$), in Y ($R^2Y$) and with cross-validated predictive power ($Q^2$). $Q^2$ values smaller than 0.098 indicate that the model has no predictive power and those greater than 0.5 indicate high predictive power. Acute angles denote positive associations, obtuse angles negative, and right angles negligible relationships.

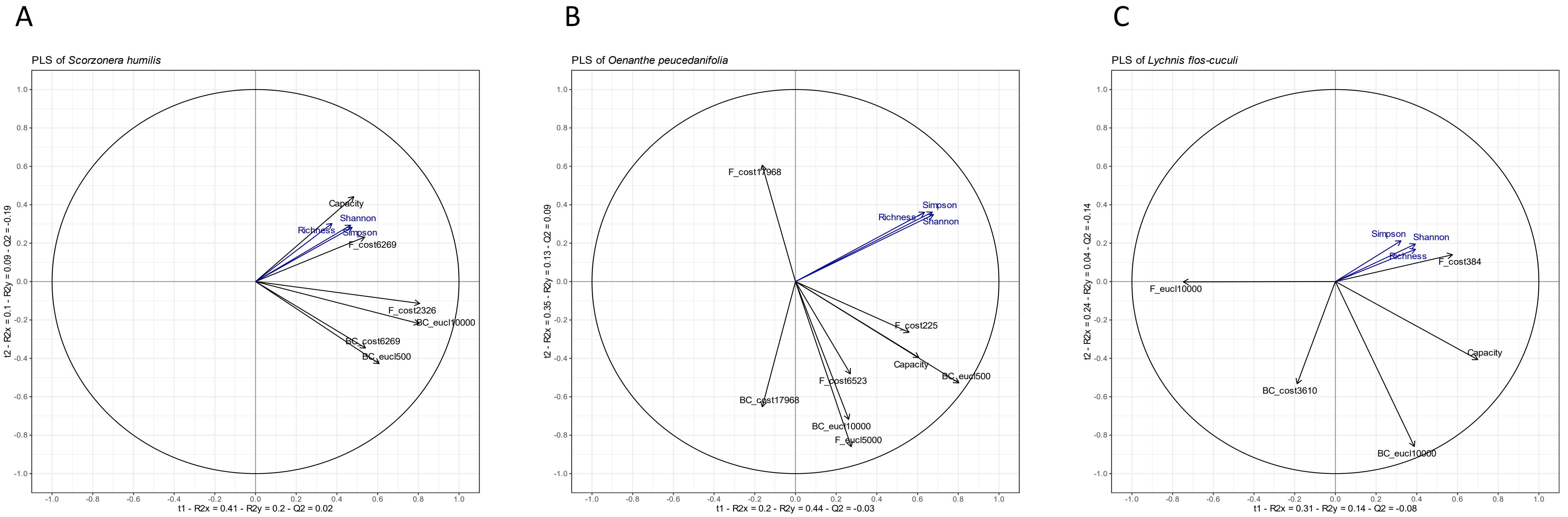

### Figure S12.

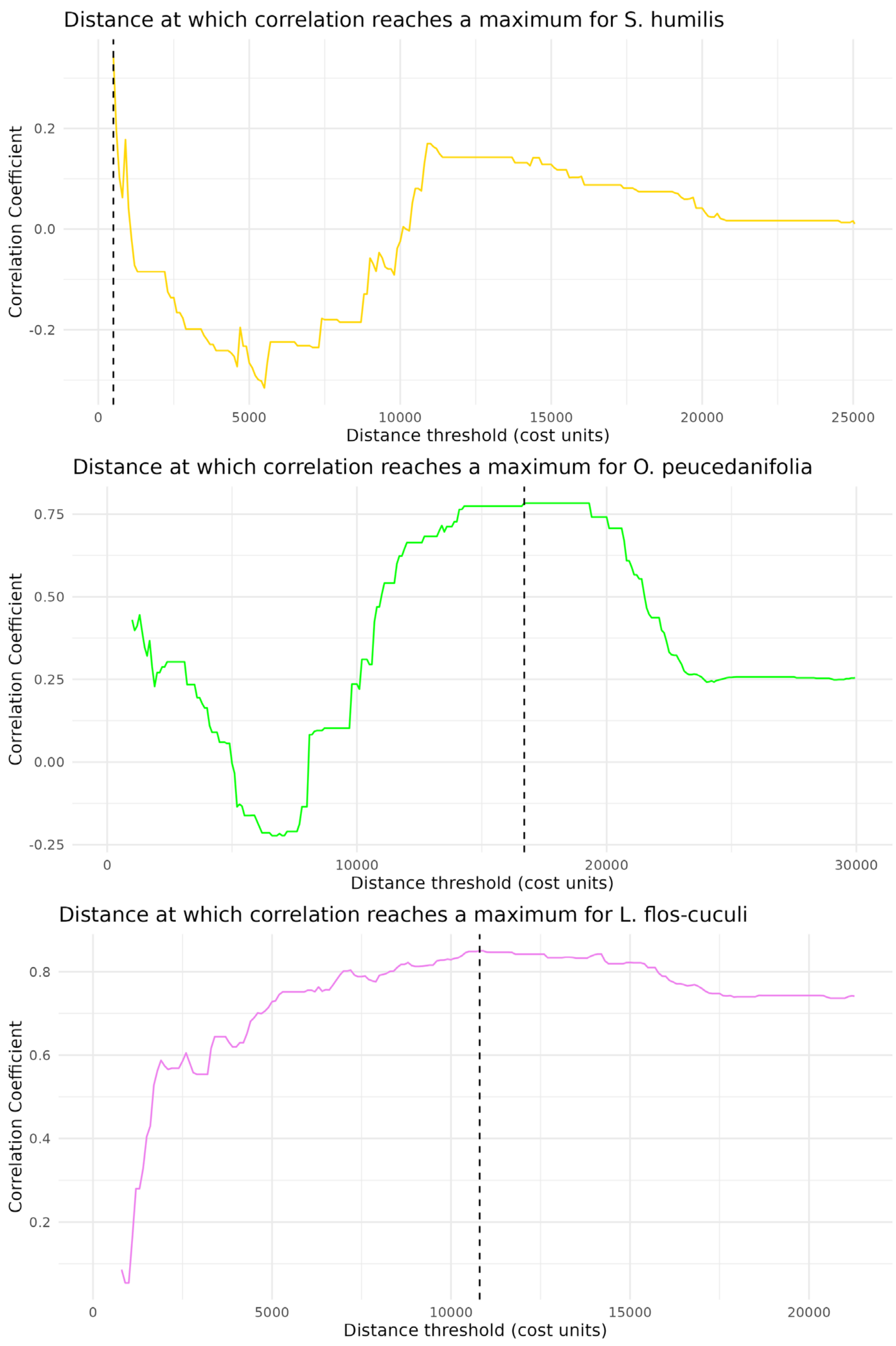


Distance of maximum correlation (DMC) depicting the distance threshold that maximizes the correlation between genetic differentiation in 3 species and the accumulated cost distance of the least cost-path. (orange=*S. humilis*, Green=*O. peucedanIfolia*, Pink=*L. flos-cuculi.* Dashed lines show the DMC for all species (*S.humilis*=500 cost units, *O.peucedanifolia*=16700, *L. flos-cuculi*=10800). This analysis reflects the distance of effective dispersal beyond which distance decreases genetic relatedness among individuals.

Figure S13.

**Generalized Dissimilarity Model (GDM) of genetic differentiation in *Oenanthe peucedanifolia*.**

I-spline functions showing the ecological distance associated with geographic distance, environmental variables, habitat patch capacity, species composition, and abundance. The y-axis represents the contribution of each predictor to genetic turnover (linearized $F_{ST}$), while the x-axis represents the gradient of each predictor. The maximum height of each curve reflects total explained turnover along that gradient, and the slope indicates the rate of allele frequency change. Significant predictors ($p < 0.05$, 999 permutations) include geographic distance (p=0.006, Predictor importance of 0.869), and the abundances of *Silaum silaus (Silsil, p=0.003, predictor importance of 3.63) Myosotis scorpioides (Myodis, p=0.015, predictor importance of 3.718)*, and shade (p=0.049, predictor importance of 10.1) indicating strong contributions of habitat structure, spatial distance, and a certain community composition to genetic differentiation.

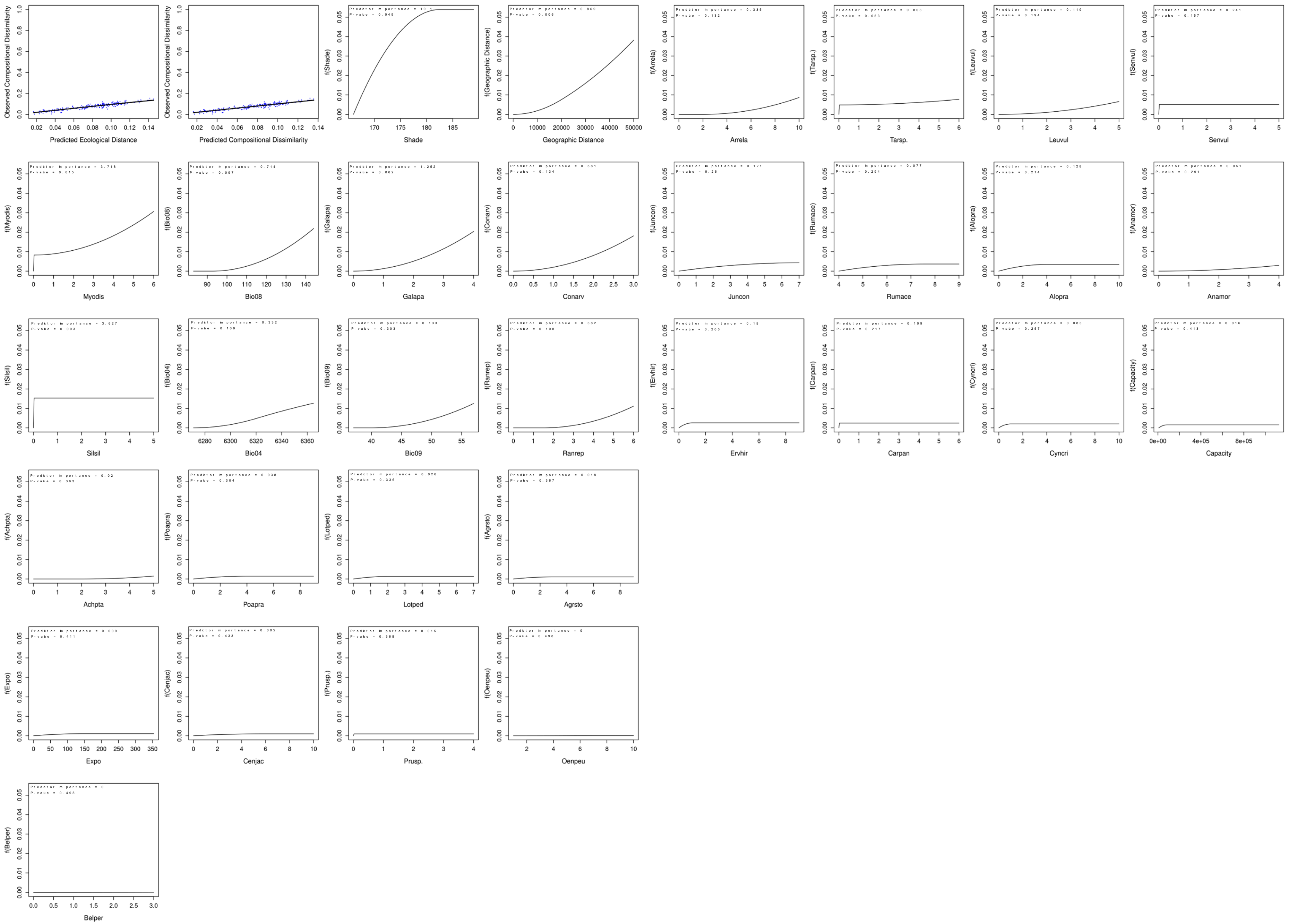

Figure S14.

#### Generalized Dissimilarity Model (GDM) of genetic differentiation in *Lychnis flos-cuculi* .

I-spline functions showing the ecological distance associated with geographic distance, environmental variables, habitat patch capacity, species composition, and abundance. The y-axis represents the contribution of each predictor to genetic turnover (linearized $F_{ST}$), while the x-axis represents the gradient of each predictor. The maximum height of each curve reflects total explained turnover along that gradient, and the slope indicates the rate of allele frequency change. Bio08 – temperature of the wettest quarter- (p = 0.001, predictor importance of 1.951 with a steep increase), and Bio18 – precipitation of the warmest quarter- (p = 0.047, predictor importance of 0.541) along with topographic slope (p = 0.049, predictor importance of 1.026), tall fescue - *Schedonorus arundinaceus* (p = 0.049, predictor importance of 0.783)- and world caraway - *Trocdaris verticillatum* (p = 0.050, predictor importance of 0.53).

**General Dissimilarity modelling of biotic and abiotic factors in *L. flos-cuculi***

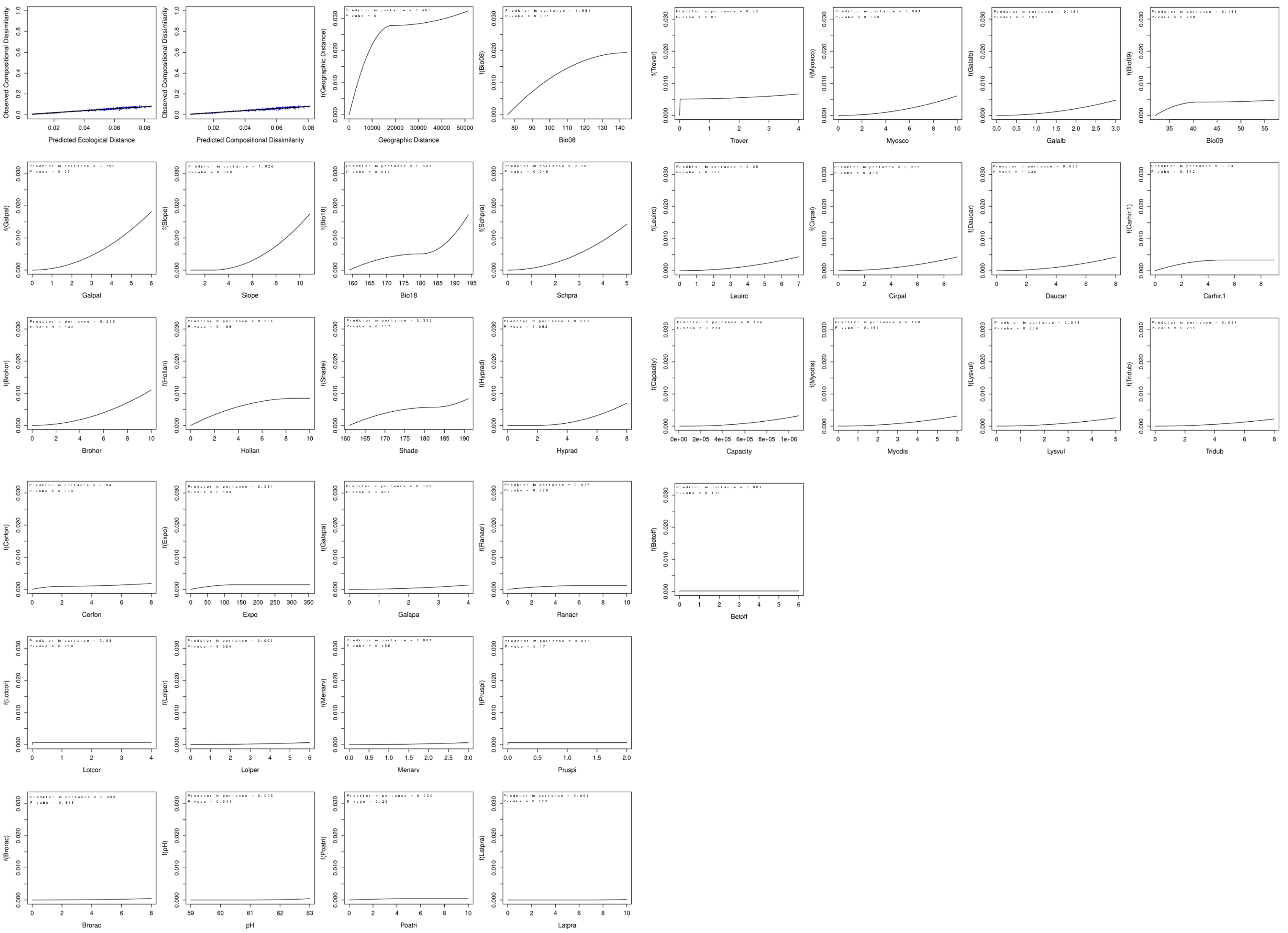

**Table S1:** Number of sampled and genotyped plants per species and per plateau. Genome sizes are given in picogram per diploid genome. Genome sizes were determined by Plant Cytometry Services (Didam, NL).

| Plateau/species | *Scorzonera humilis* | *Oenanthe Peucedanifolia* | *Lychnis flos-cuculi* | Total |
|---|---|---|---|---|
| Mornantais | 153 | 144 | 136 | 433 |
| Pélussinois | 147 | 62 | 140 | 349 |
| Annonéen | 150 | 140 | 145 | 435 |
| Total sampled | 450 | 346 | 421 | 1217 |
| **Total genotyped** | **286** | **252** | **331** | **869** |
| | 63% | 72% | 78% | |
| Genome size | 4.8pg/2c | 46.8pg/2c | 21.1pg/2c | |

**Table S2**

Environmental variables used for the species distribution modelling.

| Category | Variables | | Source |
|---|---|---|---|
| **Topography - Pedology** | Exposure – Slope – Altitude – Shading – pH - Geology | | MNT(CRAIG) SoilGrids (Hengl et al., 2014) |
| **Land use** | Agriculture – Land use – Distance to waterways – Distance to roads | | RPG (2018) / BDforests / BDtopo (https://www.ign.fr/) |
| **Climate** | Bio1=mean annual air temperature<br>Bio2=mean diurnal air temperature range<br>Bio3 =isothermality<br>Bio4=temperature seasonality<br>Bio5=mean daily maximum air temperature of the warmest month<br>Bio6=mean daily minimum air temperature of the coldest month<br>Bio7=annual range of air temperature<br>Bio8= mean daily mean air temperatures of the wettest quarter<br>Bio9= mean daily mean air temperatures of the driest quarter<br>Bio10=mean daily mean air temperatures of the warmest quarter | Bio11=mean daily mean air temperatures of the coldest quarter<br>Bio12=annual precipitation amount<br>Bio13=precipitation amount of the wettest month<br>Bio14=precipitation amount of the driest month<br>Bio15=precipitation seasonality<br>Bio16=mean monthly precipitation amount of the wettest quarter<br>Bio17=mean monthly precipitation amount of the driest quarter<br>Bio18= mean monthly precipitation amount of the warmest quarter<br>Bio19= mean monthly precipitation amount of the coldest quarter | Chelsa climate V1.2 (Karger et al.,2017) |


1. Hengl T, de Jesus JM, MacMillan RA, Batjes NH, Heuvelink GBM, et al. (2014) SoilGrids1km — Global Soil Information Based on Automated Mapping. PLOS ONE 9(8): e105992. https://doi.org/10.1371/journal.pone.0105992
2. Karger, D., Conrad, O., Böhner, J. *et al.* Climatologies at high resolution for the earth's land surface areas. *Sci Data* **4**, 170122 (2017). https://doi.org/10.1038/sdata.2017.122

#### Table S4

Nucleotide diversity (π) estimated from all sites (variant and invariant) using pixy v2.0.0.beta14 (Korunes & Samuk 2021) for each population of the three focal species. n: number of individuals; Total sites: number of genomic sites assessed; Windows: number of 10 kb windows used for estimation. In grey are populations with n ≤ 2 individuals that were excluded from average species-level diversity summaries.

| | Population | N | π (Pixy) | Total Sites | Windows |
|---|---|---|---|---|---|
| ***L. flos-Cuculi*** | a_1 | 13 | 0.0387 | 507,256 | 8959 |
| | a_10 | 13 | 0.0389 | 507,256 | 8959 |
| | a_2 | 12 | 0.0374 | 507,256 | 8959 |
| | a_3 | 12 | 0.0373 | 507,256 | 8959 |
| | a_4 | 13 | 0.0368 | 507,256 | 8959 |
| | a_5 | 13 | 0.0372 | 507,256 | 8959 |
| | a_6 | 15 | 0.0370 | 507,256 | 8959 |
| | a_7 | 14 | 0.0387 | 507,256 | 8959 |
| | a_8 | 13 | 0.0374 | 507,256 | 8959 |
| | a_9 | 14 | 0.0386 | 507,256 | 8959 |
| | m_1 | 11 | 0.0392 | 507,256 | 8959 |
| | m_10 | 15 | 0.0397 | 507,256 | 8959 |
| | m_2 | 13 | 0.0394 | 507,256 | 8959 |
| | m_3 | 12 | 0.0398 | 507,256 | 8959 |
| | m_4 | 13 | 0.0391 | 507,256 | 8959 |
| | m_5 | 9 | 0.0463 | 507,234 | 8959 |
| | m_6 | 13 | 0.0394 | 507,256 | 8959 |
| | m_7 | 6 | 0.0417 | 507,229 | 8958 |
| | m_8 | 13 | 0.0387 | 507,256 | 8959 |
| | m_9 | 13 | 0.0382 | 507,256 | 8959 |
| | p_1 | 7 | 0.0454 | 507,042 | 8957 |
| | p_10 | 13 | 0.0396 | 507,229 | 8958 |
| | p_2 | 15 | 0.0387 | 507,256 | 8959 |
| | p_3 | 13 | 0.0376 | 507,256 | 8959 |
| | p_4 | 14 | 0.0373 | 507,256 | 8959 |
| | p_5 | 13 | 0.0391 | 507,256 | 8959 |
| | p_6 | 12 | 0.0387 | 507,256 | 8959 |
| | p_7 | 11 | 0.0412 | 507,256 | 8959 |
| | p_8 | 11 | 0.0397 | 507,256 | 8959 |
| | p_9 | 12 | 0.0399 | 507,256 | 8959 |
| ***S. humilis*** | a_1 | 12 | 0.0352 | 134,894 | 4057 |
| | a_10 | 13 | 0.0349 | 134,894 | 4057 |
| | a_2 | 12 | 0.0356 | 134,894 | 4057 |
| | a_3 | 12 | 0.0355 | 134,894 | 4057 |
| | a_4 | 14 | 0.0354 | 134,894 | 4057 |
| | a_5 | 14 | 0.0350 | 134,894 | 4057 |
| | a_6 | 14 | 0.0348 | 134,894 | 4057 |
| | a_7 | 10 | 0.0355 | 134,894 | 4057 |
| | a_8 | 12 | 0.0354 | 134,894 | 4057 |
| | a_9 | 12 | 0.0343 | 134,894 | 4057 |
| | m_1 | 11 | 0.0347 | 134,894 | 4057 |
| | m_10 | 3 | 0.0378 | 134,757 | 4057 |
| | m_11 | 1 | 0.0540 | 134,021 | 4047 |
| | m_12 | 1 | 0.0505 | 111,398 | 3784 |
| | m_2 | 13 | 0.0351 | 134,894 | 4057 |
| | m_3 | 12 | 0.0350 | 134,894 | 4057 |
| | m_4 | 6 | 0.0376 | 134,894 | 4057 |
| | m_5 | 8 | 0.0356 | 134,894 | 4057 |
| | m_6 | 9 | 0.0343 | 134,894 | 4057 |
| | m_7 | 9 | 0.0351 | 134,894 | 4057 |
| | m_8 | 4 | 0.0360 | 134,877 | 4057 |
| | m_9 | 6 | 0.0360 | 134,894 | 4057 |
| | p_1 | 13 | 0.0343 | 134,894 | 4057 |
| | p_10 | 2 | 0.0405 | 129,968 | 3968 |
| | p_11 | 1 | 0.0513 | 127,763 | 3891 |
| | p_2 | 14 | 0.0351 | 134,894 | 4057 |

| | | | | | |
|---|---|---|---|---|---|
| | p_3 | 13 | 0.0354 | 134,894 | 4057 |
| | p_4 | 13 | 0.0355 | 134,894 | 4057 |
| | p_5 | 12 | 0.0356 | 134,894 | 4057 |
| | p_6 | 12 | 0.0352 | 134,894 | 4057 |
| | p_7 | 12 | 0.0353 | 134,894 | 4057 |
| | p_8 | 11 | 0.0356 | 134,894 | 4057 |
| | p_9 | 1 | 0.0430 | 52,516 | 1988 |
| ***O. peucedanifolia*** | a_1 | 11 | 0.0148 | 468,176 | 7480 |
| | a_10 | 11 | 0.0148 | 468,176 | 7480 |
| | a_2 | 13 | 0.0154 | 468,176 | 7480 |
| | a_3 | 9 | 0.0149 | 468,176 | 7480 |
| | a_4 | 9 | 0.0148 | 468,176 | 7480 |
| | a_5 | 9 | 0.0151 | 468,176 | 7480 |
| | a_6 | 7 | 0.0143 | 468,176 | 7480 |
| | a_7 | 9 | 0.0147 | 468,176 | 7480 |
| | a_8 | 11 | 0.0151 | 468,176 | 7480 |
| | a_9 | 13 | 0.0144 | 468,176 | 7480 |
| | m_1 | 12 | 0.0170 | 468,176 | 7480 |
| | m_10 | 11 | 0.0149 | 468,176 | 7480 |
| | m_2 | 15 | 0.0176 | 468,176 | 7480 |
| | m_3 | 11 | 0.0169 | 468,176 | 7480 |
| | m_4 | 10 | 0.0159 | 468,176 | 7480 |
| | m_5 | 9 | 0.0165 | 468,176 | 7480 |
| | m_6 | 11 | 0.0168 | 468,176 | 7480 |
| | m_7 | 9 | 0.0170 | 468,173 | 7480 |
| | m_8 | 9 | 0.0166 | 468,176 | 7480 |
| | m_9 | 10 | 0.0157 | 468,176 | 7480 |
| | p_1 | 11 | 0.0144 | 468,176 | 7480 |
| | p_2 | 12 | 0.0144 | 468,176 | 7480 |
| | p_3 | 12 | 0.0146 | 468,176 | 7480 |
| | p_4 | 10 | 0.0135 | 468,176 | 7480 |
| | p_5 | 2 | 0.0157 | 439,016 | 7217 |
